# An early evolutionary stage of mutualistic endosymbiosis reveals a parasitic aspect of endosymbionts

**DOI:** 10.1101/2025.05.12.653378

**Authors:** Md Mostafa Kamal, Yu-Hsuan Cheng, Li-Wen Chu, Phuong-Thao Nguyen, Chien-Fu Jeff Liu, Chia-Wei Liao, Thomas Posch, Jun-Yi Leu

## Abstract

Mutualistic endosymbiosis is a cornerstone of evolutionary innovation, enabling organisms to exploit diverse niches unavailable to individual species. However, our knowledge about the early evolutionary stage of this relationship remains limited. The association between the ciliate *Tetrahymena utriculariae* and its algal endosymbiont *Micractinium tetrahymenae* indicates an incipient stage of endosymbiosis. Although *T. utriculariae* cells rely on endosymbiotic algae to grow in low-oxygen conditions, they gradually lose the endosymbionts in aerobic conditions. Our comparative genomics reveals that the mitochondria-related genes of *T. utriculariae* are fast-evolving. Symbiotic cells display elongated mitochondria, which interact intimately with endosymbionts. Moreover, inhibiting mitochondrial fatty acid oxidation reduces host fitness but increases the endosymbiont population. Time-series transcriptomics reveal physiological fine-tuning of the host during day and night, underscoring adaptations for nutrient exchange and regulation. Notably, endosymbiotic algae downregulate photosynthesis-related genes compared with free-living cells, correlated with a substantially reduced chlorophyll content. These findings indicate that the endosymbionts exploit host metabolites to supplement reduced photosynthesis. Consistently, symbiotic *Tetrahymena* cells exhibit lower fitness under aerobic conditions than aposymbiotic cells. Our results support that mutualistic and parasitic relationships between symbiotic organisms are condition-dependent, especially at an early evolutionary stage.

## Introduction

Endosymbiosis, the intimate association by which one organism lives within the cells of another, has been a driving force in the evolution of complex life on Earth. The most profound examples are the origins of mitochondria and chloroplasts from free-living bacteria, events that gave rise to eukaryotic cells and enabled the diversification of plants and animals [1, 2]. These organelles are indispensable for energy production and photosynthesis, highlighting the evolutionary significance of endosymbiotic relationships.

Photoendosymbiosis, a specific type of endosymbiosis involving a photosynthetic organism residing within a non-photosynthetic host, is widespread across various taxa. This relationship is fundamental in marine ecosystems, particularly in coral reefs, where cnidarians harbor photosynthetic dinoflagellates of the genus *Symbiodinium*. The endosymbionts provide the host with organic compounds produced via photosynthesis, while the host offers protection and access to light and inorganic nutrients [3, 4]. Similar mutualistic associations are observed in freshwater environments, such as the endosymbiosis between the ciliate *Paramecium bursaria* and green algae of the genus *Chlorella*. In this case, the algae contribute photosynthetically-derived oxygen and nutrients, such as maltose, and in return, they receive shelter and carbon dioxide from the host [5, 6].

These well-established photoendosymbiotic relationships are characterized by a high degree of metabolic integration and mutual dependence, arising from long-term coevolution [7, 8]. The genomes of the host and endosymbiont often exhibit signs of reciprocal gene transfer and functional specialization, facilitating efficient nutrient exchange and adaptation to specific ecological niches [9, 10]. When an endosymbiotic relationship is disrupted, the fitness of both the host and endosymbiont is often compromised. However, the initial stages of endosymbiosis, i.e., when the interactions between host and endosymbiont are less integrated, remain poorly understood. Investigating nascent or early-stage symbiotic relationships is crucial to elucidate the mechanisms and selective pressures that drive the evolution of stable mutualisms [11, 12].

The ciliate *Tetrahymena utriculariae* and its algal endosymbiont *Micractinium tetrahymenae* represent a unique system for studying early-stage endosymbiosis. *T. utriculariae* is a freshwater protozoan originally found in the traps of a carnivorous plant, *Utricularia reflexa*, where it plays a role in nutrient cycling [13]. *M. tetrahymenae*, a green alga, has been identified as an endosymbiont within *T. utriculariae* [14]. The endosymbiotic algae can be transmitted vertically to progeny during cell division. However, when *T. utriculariae* cells are grown under conditions of aerobiosis and bacterial food in saturation, they gradually lose their endosymbionts [14]. This scenario contrasts with that of the *P. bursaria* system, in which endosymbionts are eliminated only when chemicals or a lack of light compromise algal growth [5, 15, 16]. Thus, the endosymbiotic relationship between *T. utriculariae* and *M. tetrahymenae* appears relatively unstable, likely representing an incipient evolutionary stage.

Environmental factors play a crucial role during the evolution of endosymbiosis. Many endosymbionts supply their hosts with unique metabolites that enable them to exploit specialized ecological niches. For instance, aphids rely on the obligate endosymbiont *Buchnera aphidicola* to synthesize essential amino acids that are absent from their phloem-sap diet, thereby allowing them to flourish in nutrient-poor environments [17]. Similarly, deep-sea clams harbor symbionts with reduced genomes that oxidize hydrogen sulfide to provide chemical energy, enabling their survival in extreme and resource-limited habitats [18]. These examples underscore how a stable and constant selective environment can promote tight host–symbiont coevolution, leading to enduring and mutually beneficial relationships—a concept comprehensively reviewed by Moya et al. [19], who elaborate on the genomic intricacies underpinning these associations. Conversely, shifts in environmental conditions can destabilize these interactions, even leading to complete symbiosis breakdown, as detailed in the work by Sachs et al. on evolutionary transitions in bacterial symbioses [20]. Such condition-dependent relationships likely occur during the early stages of symbiotic integration when the partners are not yet fully co-adapted.

*Tetrahymena* species have been studied for over a century, with *Tetrahymena thermophila* representing a model organism used in molecular and cellular biology [21, 22]. To date, *T. utriculariae* is the only *Tetrahymena* species known to form endosymbiotic relationships with green algae. The wealth of knowledge on the cell biology of *Tetrahymena* species provides an excellent resource to dissect these endosymbiotic relationships in detail.

In this study, we characterize the genomic and transcriptomic profiles of both *T. utriculariae* and *M. tetrahymenae* to identify possible pathways and physiological changes involved in their symbiotic interaction. Our phylogenetic analysis shows that mitochondria-related genes in *T. utriculariae* have undergone accelerated evolution compared to those in closely related *Tetrahymena* species. Moreover, our analyses of mitochondrial morphology and gene expression reveal specific changes in symbiotic *T. utriculariae* cells, indicating that the host mitochondria have evolved to adapt to the endosymbionts. Strikingly, many genes involved in photosynthesis are downregulated in the endosymbiotic algae, with their chlorophyll content also being reduced. This scenario raises the possibility that *M. tetrahymenae* sequesters host nutrients to support its own growth. Consistently, we observed that symbiotic *T. utriculariae* cells grew more slowly than aposymbiotic cells when cultured under aerobic conditions with bacterial food in saturation. Our data indicate that a symbiotic relationship can switch from being mutualistic to parasitic depending on the environment, providing insights into the early stages of endosymbiotic evolution.

## Results

### High-quality macronuclear genome assembly and annotation of *Tetrahymena utriculariae*

We felt that a complete genome of *T. utriculariae* was a necessary first step to understanding the genetic basis of endosymbiosis. Accordingly, we assembled a high-quality chromosome-level macronuclear (MAC) genome of *T. utriculariae* using a combination of long-read and short-read sequencing technologies (see Materials and Methods for details). The genome assembly produced a total of 181 MAC chromosomes, with a genome size of 98.0 Mb (Table 1). Annotation of the assembly yielded 23,219 genes, with protein-coding sequences averaging 2,081 base pairs (bp) in length. Despite having fewer annotated genes than two closely related species (*T. malaccensis* and *T. thermophila*), the alveolate BUSCO analysis found 97.7% of conserved genes in our assembly, similar to the BUSCO score of *T. thermophila* (97.1%, Table 1). This outcome indicates that our assembled genome captures a comprehensive set of essential genes, supporting its suitability for detailed functional and comparative genomic studies.

**Table 1.**
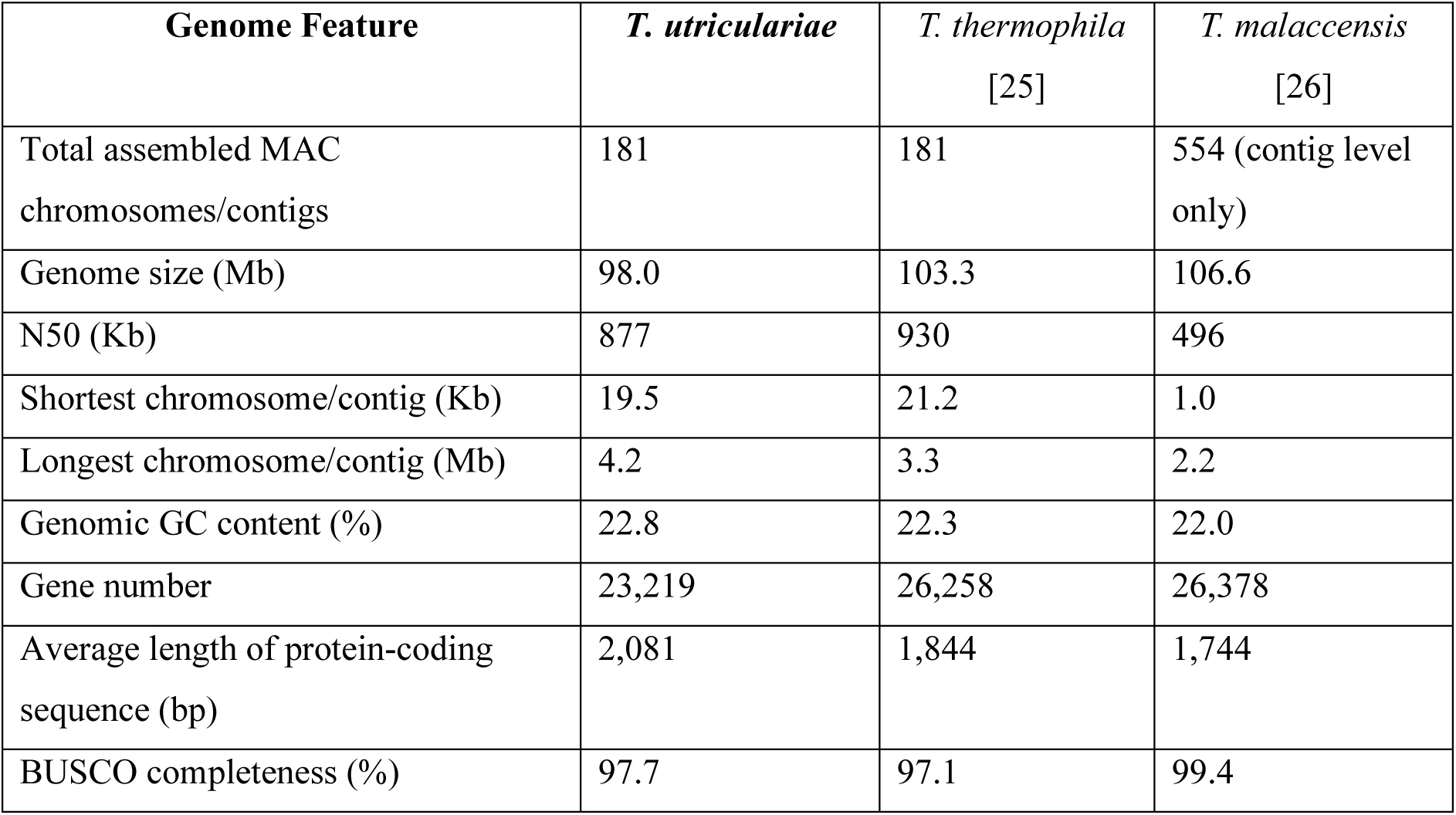
Comparative genomic attributes of *T. utriculariae*, *T. thermophila*, and *T. malaccensis*.

| Genome Feature | *T. utriculariae* | *T. thermophila*<br>[25] | *T. malaccensis*<br>[26] |
| --- | --- | --- | --- |
| Total assembled MAC chromosomes/contigs | 181 | 181 | 554 (contig level only) |
| Genome size (Mb) | 98.0 | 103.3 | 106.6 |
| N50 (Kb) | 877 | 930 | 496 |
| Shortest chromosome/contig (Kb) | 19.5 | 21.2 | 1.0 |
| Longest chromosome/contig (Mb) | 4.2 | 3.3 | 2.2 |
| Genomic GC content (%) | 22.8 | 22.3 | 22.0 |
| Gene number | 23,219 | 26,258 | 26,378 |
| Average length of protein-coding sequence (bp) | 2,081 | 1,844 | 1,744 |
| BUSCO completeness (%) | 97.7 | 97.1 | 99.4 |

A synteny analysis uncovered high collinearity (i.e., conserved gene order) between *T. utriculariae* and *T. thermophila* (Figure 1A), indicating that most genomic regions have remained relatively stable throughout their evolutionary history. However, the synteny plot also revealed genomic rearrangements. These regions of divergence may reflect species-specific adaptations. We discuss these further at the gene level in a later section.

**Figure 1.**
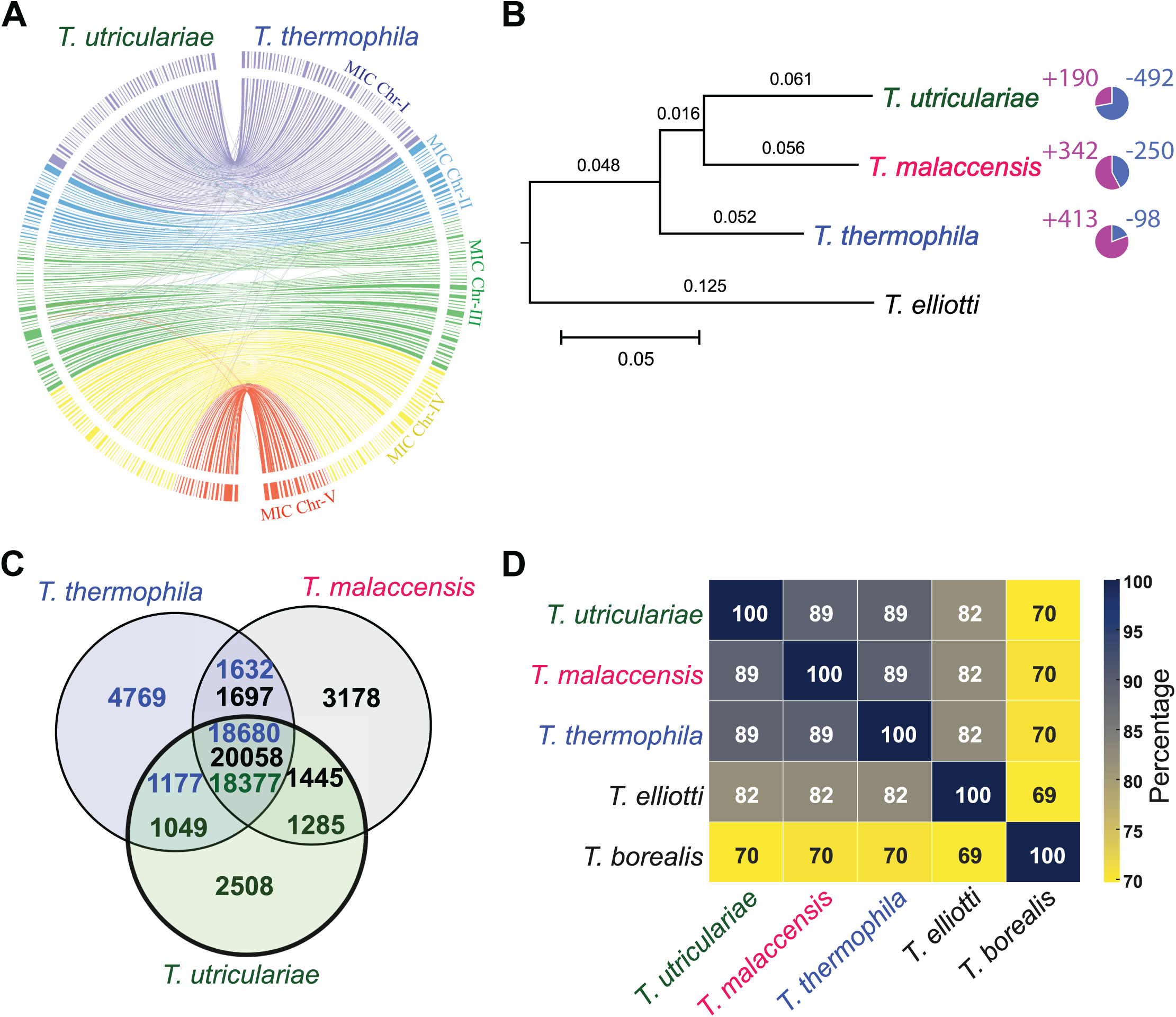
Comparative genomic analysis of *T. utriculariae* in relation to closely related ciliate species. (A) The extensive synteny we observed demonstrates considerable genomic conservation between *T. utriculariae* and *T. thermophila*. A Circos plot illustrates the syntenic relationships between the genomes of *T. utriculariae* (left) and *T. thermophila* (right). The outer rings represent macronuclear (MAC) chromosomes, color-coded according to their origin from five distinct micronuclear (MIC) chromosomes (MIC Chr I-V). Inner connecting lines represent syntenic blocks (alignments exceeding 2,000 bp), with colors corresponding to their originating micronuclear chromosomes. (B) *T. utriculariae* shares a closer evolutionary relationship with *T. malaccensis* than with *T. thermophila*, with *T. elliotti* serving as an outgroup. Maximum likelihood phylogenetic tree based on orthologous gene sequences from four *Tetrahymena* species. Branch lengths indicate genetic distance (scale bar: 0.05 substitutions per site). Pie charts represent gene family dynamics, with purple sectors indicating expanded gene families (+190, +342, and +413 for *T. utriculariae*, *T. malaccensis*, and *T. thermophila*, respectively) and blue sectors showing contracted gene families (-492, -250, and -98). (C) *T. utriculariae* shares over 79% of its genes with both *T. malaccensis* and *T. thermophila*. Venn diagram quantifying shared and unique orthologous gene clusters among *T. utriculariae* (green), *T. malaccensis* (black), and *T. thermophila* (blue). Numbers represent gene counts in each species before clustering into orthologous groups. (D) *T. utriculariae*, *T. malaccensis*, and *T. thermophila* form a closely related group with 89% amino acid identity. Average Amino Acid Identity (AAI) heatmap comparing five *Tetrahymena* species. The color gradient and numerical values indicate percentage identity between each species pair. This analysis corroborates the phylogenetic relationships and provides quantitative measures of protein-level conservation across the genus.

To clarify the evolutionary relationships between *T. utriculariae* and other species within the *Tetrahymena* genus, we used our genomics data to conduct a phylogenetic analysis on 12 *Tetrahymena* species and an outgroup, *Ichthyophthirius multifiliis*, with this latter being a close relative to Genus *Tetrahymena* [23, 24] (Supplementary Figure S1A). The resulting phylogenetic tree positioned *T. utriculariae* in a distinct clade with *T. malaccensis* and *T. thermophila*. Divergence time estimates (Supplementary Figure S1B) indicate that *T. utriculariae* and *T. malaccensis* diverged from their common ancestor with *T. thermophila* approximately 23 million years ago (MYA), and from each other around 19 MYA. To focus on this closely related clade, we reconstructed a phylogenetic tree using only *T. utriculariae*, *T. malaccensis*, and *T. thermophila*, with *T. elliotti* as an outgroup (Figure 1B), which resulted in a similar phylogenetic pattern. Despite their genetic similarity, only *T. utriculariae* has evolved into a photoendosymbiotic species, indicating that it is a species-specific adaptation.

### Comparative genomics and protein similarity between *T. utriculariae* and other *Tetrahymena* species

To explore the divergence between the genomes of *T. utriculariae* and its closely related species, we conducted an orthogroup analysis and observed that *T. utriculariae* shares >79% of its genes (18,377 out of 23,219, Supplementary Data S1A) with both *T. malaccensis* and *T. thermophila* (Figure 1C, Supplementary Data S1B). An Amino Acid Identity (AAI) analysis of 9430 one-to-one orthologous genes further reinforced the close relationship between these species. *T. utriculariae* shares 89% AAI with both *T. malaccensis* and *T. thermophila*, but the AAI declines to 82% and 70%, respectively, with the next two most closely related species, i.e., *T. elliotti* and *T. borealis* (Figure 1D).

Apart from the shared genomic content, *T. utriculariae* possesses 2,508 unique genes (∼11% of its genome, Supplementary Data S1C) not found in either *T. malaccensis* or *T. thermophila*. An examination of these *T. utriculariae*-specific genes could shed light on unique adaptations in this species. A Gene Ontology (GO) enrichment analysis of these *T. utriculariae*-specific genes indicated enrichment for biotic interactions, cellular signaling, cellular catabolism, and carbohydrate metabolism (Supplementary Figure S2, Supplementary Data S2A).

The biotic interactions and cell signaling GO categories encompass responses to other organisms and stimuli, signal transduction, and cell communication processes, which are critical for coordinating responses to internal and external stimuli [27]. Consequently, these unique genes may facilitate the ability of *T. utriculariae* to manage and respond to its symbiotic algae. The cellular catabolism and carbohydrate metabolism GO categories include autophagy, autophagosome organization, and glycogen and glucan biosynthesis, which may be involved in maintaining homeostasis and coordinating metabolic exchanges within the host-symbiont relationship [28]. Together, these *T. utriculariae*-specific genes may contribute to adaptations linked to *T. utriculariae*’s specialized photoendosymbiotic relationship with *M. tetrahymenae*.

Interestingly, among the 2,508 *T. utriculariae*-specific genes, 716 are located in non-syntenic regions compared to 1,060 non-specific genes (Supplementary Table S1, see Materials and Methods for details), revealing a significant association between gene specificity and presence of syntenic blocks (χ² = 1735.33, p < 0.001). Non-syntenic blocks typically occur near telomeres or regions hosting chromosomal rearrangements (examples are shown in Supplementary Figure S3). Notably, telomeric regions are prone to rapid genomic adaptation and restructuring [29, 30], and large-scale chromosomal rearrangements can drive evolutionary change and promote new gene formation [31, 32].

Examining the gene family expansions and contractions provided further insights into the evolutionary pressures that have shaped the *T. utriculariae* genome. The species has undergone 190 gene family expansions and 492 contractions (Figure 1B, pie charts, Supplementary Data S2B and S2C), supporting both the development of new functional capabilities and the streamlining of certain genomic aspects. The expanded gene families of *T. utriculariae* are enriched in processes related to biotic interactions, organelle organization, ion transport and homeostasis, nucleotide metabolism, protein modification, and stress responses (Supplementary Figure S2, Supplementary Data S2D). These expansions plausibly reflect host adaptations to life inside *Utricularia* traps and to hosting photosynthetic endosymbionts. For example, the enrichment of diverse membrane transporters (e.g. nutrient exporters and importers) and ion homeostasis genes implies enhanced exchange and ionic regulation between host and symbiont or prey. This scenario is consistent with those of other protist– algal symbioses. For instance, *Mesodinium rubrum* and its cryptophyte endosymbiont massively co-express nutrient transporters (such as ammonium transporters) to shuttle N^-^ sources between partners [33]. These gene family expansions may facilitate nutrient exchange with the endosymbiont and help the host cope with the stress of low oxygen levels. Similar genomic adaptations have been observed in other organisms inhabiting anoxic environments, such as certain anaerobic protists and bacteria [34, 35], which have developed specialized ion transport systems to cope with stress conditions. Additionally, organelle organization and stress responses are critical for maintaining cellular homeostasis under stress conditions or when managing the metabolic demands of symbiosis [36]. The enrichment of these processes in *T. utriculariae* suggests that this species has evolved mechanisms to regulate acidic microenvironments within its cellular compartments, such as the symbiosome that harbors the endosymbionts [36]. The expansions of genes linked to biotic interactions, ion transport, protein modification, and stress response pathways highlight *T. utriculariae*’s need for metabolic flexibility and robust cellular regulation, likely due to the challenges of maintaining a stable symbiotic relationship.

Conversely, *T. utriculariae* experienced 492 gene family contraction events. Interestingly, *T. utriculariae* has completely lost 324 of these 492 contracted gene families, meaning that the species no longer hosts any genes of those gene families. However, these same gene families are present in *T. thermophila* and *T. malaccensis*. Functional analysis of the contracted gene families revealed enrichment for GO terms associated with developmental and reproductive processes (Supplementary Figure S2, Supplementary Data S2E). Notably, “meiosis I” and “homologous recombination” (cell cycle/DNA metabolism categories) and “cell adhesion” (cell signaling category) are overrepresented among the contracted gene families. The loss of meiosis-related genes potentially indicates that *T. utriculariae* relies on a streamlined set of core proteins for sexual reproduction or displays reduced sexual reproduction. Similarly, having fewer cell adhesion genes may lead to a reduction in cell-cell interactions, which could possibly be involved in cell mating. Gene family contractions in core processes can reflect evolutionary specialization. For instance, other studies have shown that loss of redundant genes in essential pathways can accompany niche adaptation and speciation [37]. Thus, the contracted GO categories imply that *T. utriculariae* has pared down redundancy in certain fundamental pathways (e.g. sexual cell division and adhesion) for lineage-specific adaptation.

### *T. utriculariae* mitochondrial genome evolution reflects its adaptations to unique ecological and symbiotic pressures

We successfully assembled the mitochondrial genome of *T. utriculariae* into a linear structure, with a total size of 51,725 bp, which is slightly larger than the respective genomes of related species such as *T. thermophila* and *T. malaccensis* (Figure 2A and Supplementary Table S2). The gene content includes 44 protein-coding genes, 7 tRNA genes, and 6 rRNA genes, with a notably larger intergenic region (6534 bp) compared to its relatives and an A+T content of 78.54%.

**Figure 2:**
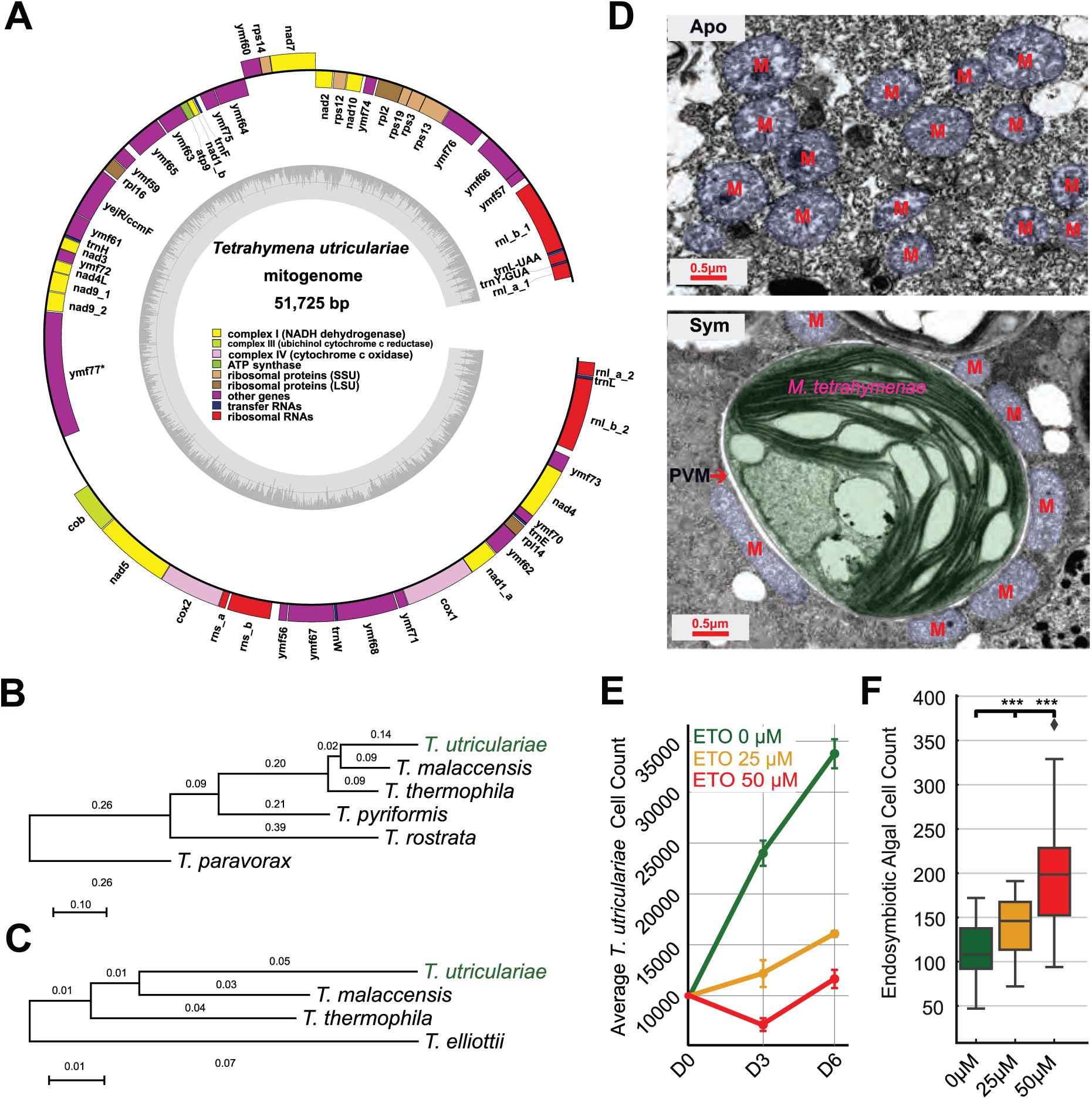
Mitochondria of *T. utriculariae* display a unique evolutionary trajectory specific to its symbiotic lifestyle. (A) Map of the *Tetrahymena utriculariae* mitochondrial genome (51,725 bp); for presentation purposes, the linear genome has been rounded. The genome encodes 44 protein-coding genes, 7 tRNAs, and 6 rRNAs. Genes are color-coded according to their respective functions, including complex I NADH dehydrogenase (yellow), complex III ubiquinol cytochrome c reductase (light brown), complex IV cytochrome c oxidase (pink), ATP synthase (light green), ribosomal proteins (brown; SSU and LSU), other genes (purple), transfer RNAs (blue), and ribosomal RNAs (red). The gene ymf77* is marked with an asterisk to indicate that it contains introns, a notable feature of this mitochondrial genome. The inner gray circle represents GC content variation across the genome. (B) Mitochondrial DNA (mtDNA)-encoded genes (n = 44) in *T. utriculariae* change more quickly than those of the other two closely related species. Maximum likelihood phylogenetic tree constructed from mitochondrial-encoded genes across six *Tetrahymena* species. Branch lengths indicate genetic distance (scale bar: 0.10 substitutions per site). The phylogenetic relationships between *T. utriculariae*, *T. malaccensis*, and *T. thermophila* are consistent with those determined from the whole-genome tree. (C) Nucleus-encoded mitochondrial genes (n = 518) in *T. utriculariae* change more quickly than those of the other two closely related species. Maximum likelihood phylogenetic tree derived from nucleus-encoded mitochondrial genes from four *Tetrahymena* species. Scale bar represents 0.01 substitutions per site. (D) The morphology of mitochondria in symbiotic cells differs from those of aposymbiotic cells. Transmission electron micrographs comparing the mitochondrial morphology of aposymbiotic (Apo, upper panel) and symbiotic (Sym, lower panel) *T. utriculariae* cells. Mitochondria (M, highlighted in blue pseudocolor) in aposymbiotic cells display a typical rounded morphology. In contrast, the mitochondria of symbiotic cells exhibit elongated structures that closely associate with and wrap around the perialgal vacuole membrane (PVM, red arrow) containing the endosymbiotic *M. tetrahymenae* alga (green pseudocolor). Scale bars: 0.5 μm. See Supplementary Figure S4 for additional images. (E) Inhibiting mitochondrial fatty acid oxidation compromises *T. utriculariae* growth. Cells were treated with the carnitine palmitoyltransferase-1 inhibitor Etomoxir (ETO) at two concentrations (25 μM and 50 μM) and monitored over 6 days. Control cells (0 μM) showed robust exponential growth, whereas treatment with 25 μM or 50 μM ETO resulted in dose-dependent growth inhibition. Error bars represent standard error of the mean from three biological replicates. (F) Inhibiting mitochondrial fatty acid oxidation increases endosymbiotic algal numbers. Endosymbiotic algal abundance in *T. utriculariae* was quantified after 6 days of ETO treatment. Box plots show a significant dose-dependent increase in endosymbiont numbers with increasing ETO concentration (0 μM, 25 μM, and 50 μM). Fifty cells were counted in each sample. Boxes represent the interquartile range with median line; whiskers extend to minimum and maximum values within the 1.5× interquartile range. ***; p < 0.001 (one-way ANOVA followed by Tukey’s post-hoc test).

We constructed a maximum likelihood tree to perform phylogenetic analysis on the mitochondrial-encoded genes (Figure 2B). The phylogenetic relationships between *T. utriculariae*, *T. malaccensis*, and *T. thermophila* based on mitochondrial genes are consistent with those reflected in the whole-genome tree (Figure 1B). However, we noticed that the branch for *T. utriculariae* is much longer than those of *T. malaccensis* and *T. thermophila*, indicating that the mitochondrial DNA (mtDNA)-encoded genes in *T. utriculariae* change more quickly. mtDNA-encoded proteins often interact with nuclear-encoded mitochondrial proteins to execute their functions. Accordingly, we assessed if the nuclear-encoded mitochondrial proteins (see Methods for nuclear-encoded mito-protein annotation) co-evolved to exhibit a similar pattern. Indeed, we determined that the nuclear-encoded mitochondrial genes in *T. utriculariae* also change more quickly than those of *T. malaccensis* and *T. thermophila* (Figure 2C). These data indicate that the mitochondrial genome of *T. utriculariae* is subjected to unique evolutionary pressures.

We performed transmission electron microscopy (TEM) on *T. utriculariae* mitochondria and, interestingly, we observed that they are remodeled in symbiotic cells. Aposymbiotic *T. utriculariae* cells predominantly hosted rounded mitochondria (Figure 2D and Supplementary Figure S4A). In contrast, the mitochondria of symbiotic cells were elongated and intimately associated with the membrane of the perialgal vacuole (PV) that houses the endosymbiont, with most non-elongated mitochondria not in contact with the PV membrane (Figure 2D, Supplementary Figure S4B and S4C). The elongated shape and close proximity of mitochondria to the PV may facilitate efficient energy transfer, metabolite exchange, and redox homeostasis at the host-symbiont interface. Alternatively, as shown in a recent study on parasitic *Toxoplasma gondii*, mitochondrial wrapping could sequester host fatty acids to limit nutrient availability to the parasite and control its proliferation [38]. This scenario raises the possibility that mitochondrial remodeling in *T. utriculariae* may also contribute to regulating the endosymbiont population.

To test the regulatory role of fatty acid oxidation and mitochondria in endosymbiosis, we treated symbiotic cells with etomoxir to inhibit fatty acid oxidation in mitochondria. Etomoxir is an irreversible inhibitor of carnitine palmitoyltransferase 1 (CPT1), a rate-limiting enzyme in mitochondrial fatty acid oxidation [39]. Etomoxir treatment of symbiotic *T. utriculariae* cells resulted in a dose-dependent reduction in host growth rates (Figure 2E), supporting that mitochondrial fatty acid oxidation is crucial for sustaining host cell growth and metabolic activity. However, inhibition of host fatty acid oxidation also significantly increased the number of endosymbionts within the host cells (Figure 2F), indicating that mitochondrial fatty acid oxidation also acts as a mechanism to regulate endosymbiont numbers. This specific regulatory mechanism in *T. utriculariae* may explain the unique evolutionary pattern of its mitochondrial genes.

### Transcriptomic analysis reveals dynamic host responses to symbiosis and environmental conditions

Building upon our comprehensive genomic analysis of *T. utriculariae*, we explored the transcriptomic landscape to uncover the molecular mechanisms underlying its endosymbiotic relationship with *M. tetrahymenae*. We conducted a time-series RNA sequencing (RNA-seq) experiment comparing symbiotic (Sym) and aposymbiotic (Apo) *T. utriculariae* cells at six time-points (AM8, AM11, PM2, PM5, PM8, PM11) through a single 24-hour light-dark cycle (see Methods for details) (Supplementary Data S3A). This approach allowed us to capture temporal changes in gene expression associated with symbiosis and environmental cues.

A principal component analysis (PCA) of the transcriptomic data revealed that symbiotic cells clustered distinctly from aposymbiotic cells, indicating that the symbiotic state is the primary factor influencing gene expression patterns, accounting for 53.6% of the total variance along the first principal component (PC1) (Figure 3A). Light conditions also contributed to varied gene expression within symbiotic cells, as samples collected during light periods (AM11, PM2, PM5) were well separated from those collected in the transition to or during dark periods (AM8, PM8, PM11), underscoring the combined influence of symbiosis and light on the host’s transcriptional landscape.

**Figure 3:**
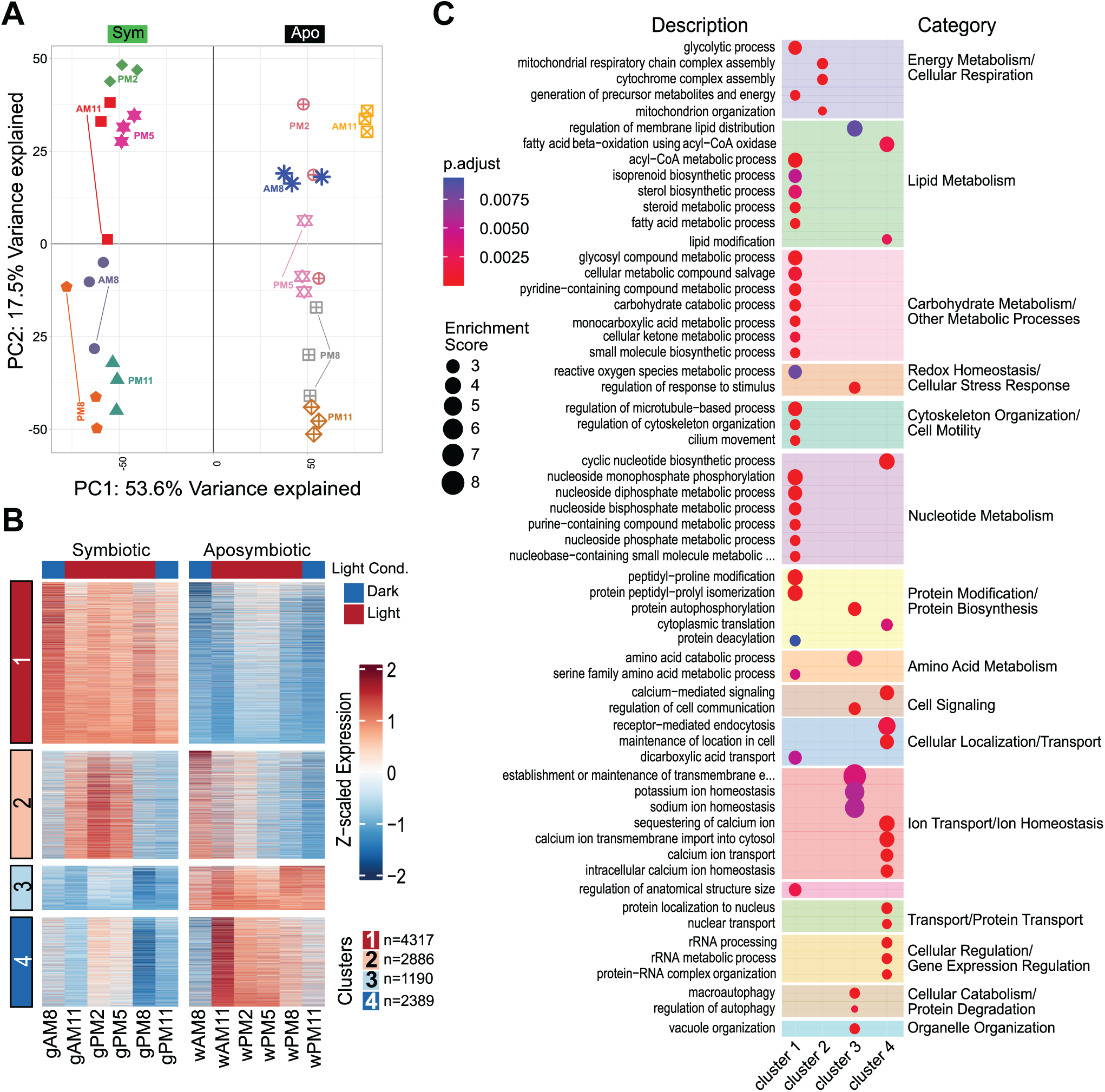
Transcriptomic dynamics of *T. utriculariae* reveal distinct metabolic signatures in symbiotic and aposymbiotic states across diurnal cycles. (A) Principal Component Analysis (PCA) of *T. utriculariae* transcriptomes at six time-points in symbiotic (Sym, solid symbols) and aposymbiotic (Apo, hollow symbols) states. PC1 (53.6% variance) distinguishes symbiotic status, whereas PC2 (17.5% variance) separates samples by time of day. (B) K-means clustering of differentially expressed genes yielded four major clusters (1-4) across experimental conditions (n=6 lines each for the symbiotic and aposymbiotic conditions). Cluster 1 (n=4,317): genes upregulated in symbiotic cells; Cluster 2 (n=2,886): genes with peak expression during midday in symbiotic cells; Cluster 3 (n=1,190): genes expressed in aposymbiotic cells; Cluster 4 (n=2,389): genes with morning expression in aposymbiotic cells. (C) Gene Ontology enrichment analysis of the four gene expression clusters. Circle size represents enrichment score, and color intensity indicates statistical significance (adjusted p-value). These enrichment patterns reveal the key metabolic and cellular processes differentially regulated in symbiotic versus aposymbiotic *T. utriculariae* throughout the diurnal cycle.

We defined a gene as being differentially expressed if expression in symbiotic cells was two-fold higher or lower than that in aposymbiotic cells (adjusted p-value < 0.01, Student’s t-test with Benjamini-Hochberg correction) at any time-point (Supplementary Data S3B). Using this cutoff, we identified 10,782 differentially expressed genes that could be roughly classified into four clusters (Figure 3B).

One prominent set of genes (Cluster 1, n = 4,317) exhibited stable expression in symbiotic cells across all time-points, but were downregulated in aposymbiotic cells (Figure 3B, Supplementary Data S3C). GO analysis revealed that these genes are heavily involved in energy, lipid and carbohydrate metabolism, homeostasis, and cytoskeleton organization (Figure 3C, Supplementary Data S3D). Genes associated with energy generation were consistently expressed, indicative of host reliance on efficient energy production, possibly to meet the metabolic demands of both the host and symbiont [40]. Given that our etomoxir treatment experiments revealed that *T. utriculariae* cells adjust fatty acid oxidation pathways to control the endosymbiont population (Figure 2F), it may explain why lipid metabolism pathways are upregulated in symbiotic cells. Furthermore, the host may actively contribute to nutritional needs of its endosymbionts, supplying key metabolites to sustain the symbiotic balance [41]. The enrichment for genes linked to cytoskeleton organization and microtubule-based movement indicates that these genes are involved in maintaining the cellular structures necessary for housing endosymbionts, potentially stabilizing the perialgal vacuole in which the algae reside [42, 43]. These core metabolic and structural processes appear to be essential for the continuous support of the endosymbiont, irrespective of external light conditions, underscoring the stable and foundational role of this gene set in symbiotic maintenance.

Cluster 2, comprising 2,886 genes, showed increased expression in symbiotic cells during midday light conditions (Figure 3B, Supplementary Data S3C). These genes are enriched in mitochondrial respiratory chain complex assembly and organization (Figure 3C, Supplementary Data S3D). Upregulation of mitochondrial genes during light exposure indicates coordination between host mitochondrial function and the endosymbiont’s photosynthetic activity [44, 45]. Together with the observation that mitochondria-related genes in *T. utriculariae* evolve more rapidly and that mitochondrial functions contribute to regulating endosymbiont numbers (Figure 2B, 2C, and 2F), this outcome illustrates the significance of host mitochondrial functions in endosymbiosis.

In contrast, the 1,190 genes of Cluster 3 were significantly upregulated in aposymbiotic cells.

GO analysis revealed enrichment for autophagy-related processes, vacuole organization, and ion homeostasis (Figure 3C, Supplementary Data S3D). The induction of autophagy indicates that aposymbiotic cells activate internal recycling mechanisms in the absence of endosymbionts, repurposing cellular components to maintain homeostasis [46, 47]. The upregulation of genes associated with vacuole organization further supports the adaptation of intracellular structures for nutrient management.

Another set of genes (Cluster 4, n = 2,389) showed high expression during early light exposure in aposymbiotic cells, emphasizing the host’s ability to respond to light cues even without the symbiont. These genes were significantly enriched in processes related to calcium ion homeostasis, nuclear transport, and rRNA metabolism. Although it remains unclear why aposymbiotic cells need to change the expression of those genes in these particular conditions, the data indicate that aposymbiotic cells are highly responsive to environmental light, adjusting their physiology to optimize cellular functions.

Collectively, these findings demonstrate that *T. utriculariae* orchestrates a complex transcriptional response to balance symbiotic maintenance with different survival strategies, which are modulated by both symbiotic state and environmental factors such as light.

### Endosymbiont *Micractinium tetrahymenae* exhibits metabolic shifts during symbiosis

To better understand the symbiotic relationship from the endosymbiont’s perspective, we sequenced and annotated the genome of *M. tetrahymenae*. This was accomplished using a hybrid assembly approach that combined Illumina paired-end short reads for accuracy and Oxford Nanopore long reads for contiguity. The assembly was polished iteratively with both data types, resulting in a high-quality genome characterized by a size of 66.4 Mb, a GC content of 65.29%, and 97.1% BUSCO completeness (Table 2). Annotation of the nuclear genome was performed by incorporating RNA-seq data to enhance gene model predictions, leading to the identification of 14,229 protein-coding genes. A one-to-one orthologous gene-based maximum likelihood phylogenetic tree clustered *M. tetrahymenae* with *M. conductrix* (Figure 4A). Additionally, we assembled the plastid and mitochondrial genomes (Supplementary Figure S5A and 5B), which displayed sizes of 133,089 bp and 91,655 bp, respectively, encoding 79 and 32 protein-coding genes (Table 2). Average protein lengths for the nuclear, plastid, and mitochondrial genomes are 478, 265, and 262 amino acids, respectively (Table 2).

**Figure 4:**
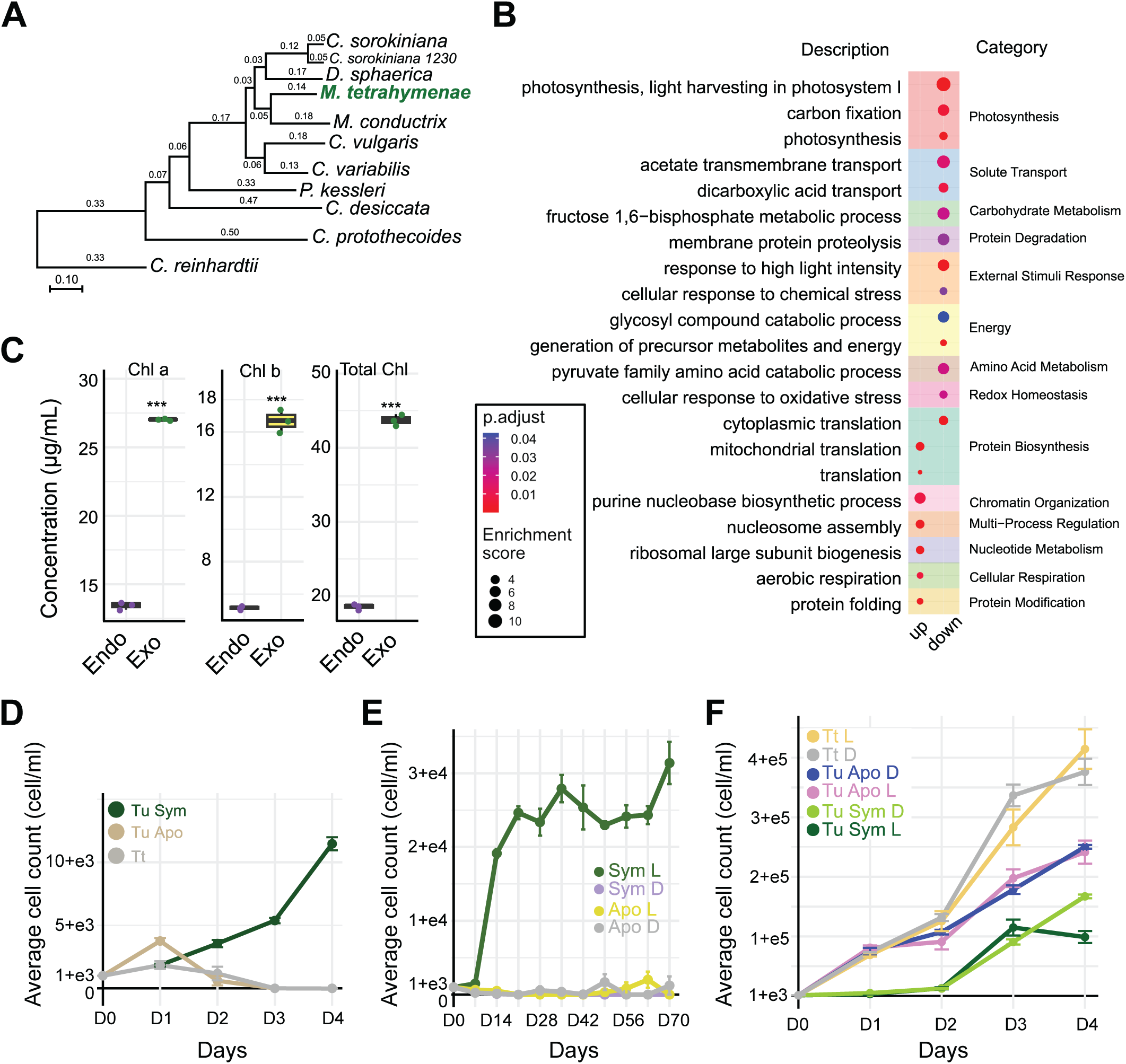
Transcriptomic and physiological responses of *M. tetrahymenae* in symbiotic and free-living states. (A) *M. tetrahymenae* shares a close relationship with *M. conductrix*, with both species being capable of establishing endosymbiotic relationships with ciliates. Maximum likelihood phylogenetic tree showing the evolutionary position of *M. tetrahymenae* (highlighted in green) among other chlorophyte algae. Branch lengths indicate genetic distance (scale bar: 0.10 substitutions per site). (B) Gene Ontology (GO) enrichment analysis of differentially expressed genes in symbiotic vs. free-living *M. tetrahymenae*. Upregulated genes in the symbiotic state are primarily involved in protein biosynthesis, cellular respiration, and nucleic acid metabolism, whereas downregulated genes are related to photosynthesis, carbon fixation, and responses to light and chemical stresses. Circle size indicates enrichment score, and color represents adjusted p-value significance. (C) Endosymbiotic algae exhibited significantly lower concentrations of chlorophyll a, chlorophyll b, and total chlorophyll than free-living algae, implying reduced photosynthetic capacity in the symbiotic state. Chlorophyll content measurements to compare symbiotic (*Endo*) vs. free-living (*Exo*) *M. tetrahymenae*. ***; p < 0.001 (Student’s t-test). (D) Symbiotic *T. utriculariae* (Tu Sym) demonstrates robust growth under hypoxic conditions, whereas aposymbiotic *T. utriculariae* (Tu Apo) and *T. thermophila* (Tt) show limited growth that plateaus or declines after the first few days. This outcome indicates that the endosymbiotic relationship provides *T. utriculariae* with a significant selective advantage in oxygen-depleted environments. Error bars represent the standard error of the mean from three biological replicates. (E) Growth curves showing cell counts of symbiotic and aposymbiotic *T. utriculariae* under hypoxic conditions. Symbiotic cells in light conditions (Sym L) exhibited robust growth, reaching ∼32,000 cells/ml by day 70, whereas symbiotic cells in constant darkness (Sym D), aposymbiotic cells in light (Apo L), and aposymbiotic cells in darkness (Apo D) all maintained minimal growth, i.e., below 1,000 cells/ml, in the same timeframe. These results demonstrate that the symbiotic relationship with algal endosymbionts and light availability are both necessary for *T. utriculariae* to thrive in oxygen-depleted environments. (F) Growth curves of *T. utriculariae* (Tu) and *T. thermophila* (Tt) under aerobic conditions and provided with bacterial food in saturation over a 4-day period. Growth is compared between symbiotic (Sym) and aposymbiotic (Apo) *T. utriculariae* under light/dark cycles (L) or constant darkness (D). *T. thermophila* cells were used as a control for sufficient food supply since they grew much faster and consumed more bacteria. Error bars represent the standard error of the mean from three biological replicates.

**Table 2:** Genomic features of the *Micractinium tetrahymenae* nuclear, plastid, and mitochondrial genomes.

| Genome feature | Nuclear genome | Plastid genome | Mitochondrial genome |
| --- | --- | --- | --- |
| Genome size (Mb) | 66.4 | 0.133 | 0.0917 |
| Total assembled scaffolds/contigs | 67 | – | – |
| N50 (Mb) | 3.29 | – | – |
| Genomic GC content (%) | 65.29 | 34.22 | 30.25 |
| Gene number | 14,229 | 79 | 32 |
| Shortest protein-coding sequence (bp) | 111 | 96 | 225 |
| Longest protein-coding sequence (bp) | 30,000 | 4,470 | 2,019 |
| Average length of protein-coding sequence (bp) | 1,434 | 794 | 787 |
| Median length of protein-coding sequence (bp) | 1,134 | 483 | 651 |
| BUSCO completeness (%) | 97.1 | – | – |

Using this high-quality genome, we analyzed the transcriptomic profiles of *M. tetrahymenae* in its endosymbiotic and free-living states, as collected at midday (2 pm) (Supplementary Data S4A). Differential expression analysis identified 1,388 genes significantly upregulated and 845 genes significantly downregulated (fold-change > 2, adjusted p-value < 0.01, Student’s t-test with Benjamini-Hochberg correction) in endosymbiotic *M. tetrahymenae* compared to free-living cells (Supplementary Data S4B). Notably, genes associated with protein biosynthesis, nucleotide biosynthesis, and aerobic respiration were upregulated in the endosymbiotic *M. tetrahymenae* (Supplementary Data S4C), indicating increased metabolic and biosynthetic activities. This shift toward enhanced metabolic processes supports that endosymbiotic *M. tetrahymenae* cells can grow vigorously inside the host.

Intriguingly, genes linked to GO terms relating to photosynthesis and light-harvesting complexes, such as light harvesting in photosystem-I (GO:0009768), as well as carbon fixation (GO:0015977), were downregulated in the endosymbiotic state (Figure 4B, Supplementary Data S4C). This outcome indicates that the endosymbiont reduces its photosynthetic activity when residing within the host. Detection of GO categories enriched for dicarboxylate and succinate transport processes (GO:0015740, GO:0006835) further indicates a metabolic shift by endosymbionts toward reduced photosynthesis (Figure 4B, Supplementary Data S4C). How can endosymbiotic algal cells support increased metabolic and biosynthetic activities when their levels of photosynthesis are decreased? Our etomoxir treatment experiments (Figure 2F) imply that the endosymbionts utilize nutrients and energy provided by the host, transitioning from autotrophic to a more heterotrophic lifestyle.

To further explore the effect of reduced photosynthetic gene expression by endosymbionts, we measured the chlorophyll content of endosymbiotic and free-living algal cells, which is a vital light-absorbing component of photosynthesis. Endosymbiotic *M. tetrahymenae* exhibited significantly lower levels of chlorophyll a, b, and total chlorophyll compared to free-living cells (Figure 4C). Together, these results indicate that the photosynthetic capabilities of endosymbionts are diminished within the host environment. It is likely then that endosymbionts take up nutrients from the host to offset their reduced generation of photosynthetic products, raising the possibility of parasitic interactions.

### Symbiosis imposes a metabolic burden on *T. utriculariae*

*T. utriculariae* was originally discovered inside the traps of *U. reflexa*, where oxygen levels are low due to the decomposition of organic matter and limited gas exchange [48, 49]. It has been proposed previously that endosymbiotic *M. tetrahymenae* provide essential oxygen to sustain host growth under this condition [14, 49]. Consequently, we cultured symbiotic and aposymbiotic *T. utriculariae* cells under artificial hypoxic conditions and monitored them for both short (4 days) and long (70 days) durations (Figure 4D and 4E). Only symbiotic cells under a light-dark cycle exhibited robust and sustained proliferation, maintaining a high cell density throughout both experimental periods. In contrast, symbiotic cells under constant dark or aposymbiotic cells showed minimal to no growth (Figure 4E), indicating that while the endosymbiont provides a survival advantage to the host under hypoxic conditions, this benefit depends entirely on light exposure.

Interestingly, we observed a different growth pattern when *T. utriculariae* cells were cultured in aerobic conditions. Aposymbiotic cells displayed higher growth rates than symbiotic cells, whether under light-dark or constant-dark conditions (Figure 4F). This result indicates that when oxygen is not limited, the endosymbiotic relationship imposes a metabolic cost on the host. More importantly, the cost cannot be alleviated by the photosynthetic activity of the endosymbionts. This outcome is consistent with our endosymbiont transcriptome data showing that *M. tetrahymenae* inside *T. utriculariae* switches from autotrophy to mixotrophy. It is likely that endosymbiotic *M. tetrahymenae* also imposes similar costs under hypoxic conditions. However, the benefit of the oxygen supply exceeds the metabolic burden, allowing symbiotic *T. utriculariae* to proliferate (Figure 4D and 4E). The parasitic aspect of endosymbiotic *M. tetrahymenae* may explain why the host needs to evolve specific mechanisms involving mitochondria and fatty acid oxidation to control the endosymbiont population (Figure 2).

Collectively, these findings indicate that the endosymbiotic association between *T. utriculariae* and *M. tetrahymenae* is highly condition-dependent. Although *M. tetrahymenae* limits host proliferation in oxygenic environments, it provides a clear survival benefit under long-term hypoxic conditions. This scenario strongly contrasts with the stable relationship between *P. bursaria* and *Chlorella spp*, in which aposymbiotic cells consistently exhibited lower fitness than symbiotic cells under all tested conditions [50]. Thus, our findings highlight the complex interplay between mutualism and parasitism at the early evolutionary stages of endosymbiosis.

## Discussions

The symbiotic relationship between the freshwater ciliate *T. utriculariae* and its algal endosymbiont *M. tetrahymenae* offers a unique model for studying the early stages of endosymbiosis. Our integrative analysis reveals that this association deviates from classical mutualistic photoendosymbiosis, instead being characterized by conditional benefits and significant fitness costs to the host. Our findings provide insights into the incipient evolutionary stage of endosymbiosis, i.e., before a stable relationship is established.

Multiple lines of evidence support that the mitochondria of *T. utriculariae* have specifically evolved to support its unique lifestyle. We observed fast-evolving patterns in mitochondria-related genes and elongated mitochondria closely associated with endosymbionts (Figure 2B-2D, and Supplementary Figure S4B). Moreover, the genes involved in mitochondrial organization and respiration are significantly upregulated in symbiotic host cells (Figure 3C). These adaptations may reflect a shift towards anaerobic metabolism or enhanced efficiency under low-oxygen conditions, as suggested previously for other anaerobic eukaryotes such as *Blastocystis* spp. and certain ciliates [51, 52]. However, our experiments reveal an additional role for mitochondrial fatty acid oxidation in regulating endosymbiont proliferation. In particular, when we reduced host mitochondrial fatty acid oxidation using etomoxir, *T. utriculariae* cells displayed decreased growth but increased numbers of endosymbionts (Figure 2E and 2F). Previous studies have shown that host mitochondria reorganize and wrap around pathogen-hosting vacuoles, in which several evolutionarily divergent prokaryotic and eukaryotic pathogens, including the bacteria *Legionella pneumophila* and *Chlamydia psitacci*, and the alveolate parasite *Toxoplasma gondii*, can proliferate [53–58]. In the case of *T. gondii*, enhanced host mitochondrial fatty acid oxidation has been demonstrated to limit the parasite’s ability to sequester fatty acids, allowing the host to control parasite number [38]. It is possible that *T. utriculariae* cells have evolved a similar mechanism to tightly regulate the endosymbiont population, thereby lowering the host metabolic burden attributable to the endosymbiotic algae.

When the current report was in preparation, Kelly et al. published a study in which they analyzed gene expansions in the *T. utriculariae* and *M. tetrahymenae* genomes [59]. Their results indicate that gene families related to stress response have been expanded in both the host and endosymbiont as their symbiotic relationship evolved. This finding further supports the notion that, at the early stage of endosymbiosis, host and endosymbiont coexistence represents a stressful environment to which both organisms need to adapt before they can become fully integrated.

Our findings underscore the importance of environmental factors in driving the evolution of endosymbiosis. Studies on other organisms inhabiting extreme environments, such as deep-sea vents or anaerobic sediments, have shown that environmental pressures can drive unique symbiotic adaptations [60]. It is possible that the ancestors of *M. tetrahymenae* escaped from the digestive system of *T. utriculariae* after they were engulfed simply due to their specific cell wall composition. However, these surviving algal cells might have quickly sensed the nutrient-rich environment within the host cells and then started exploiting those conditions. In addition, living inside a ciliate cell protects the endosymbiotic algal cells from viral infection, which is prevalent in aquatic environments [61, 62]. These parasite-like algal cells could kill the host if they proliferated without impediment, thus forcing the host cells to evolve a mechanism to control the population. Nevertheless, due to their reduced fitness, infected ciliate cells are more likely to be outcompeted by uninfected cells and thus lost from the population. However, the low-oxygen environment within *U. reflexa* traps represents a critical turning point during the evolution of the endosymbiotic relationship since only infected *T. utriculariae* cells can propagate and exploit this nutrient-rich environment. However, *Utricularia* traps constantly grow and die, meaning that infected *T. utriculariae* cells have to migrate and thrive under both hypoxic and aerobic conditions. If such an ecological condition persists for a long time, it will further select for *T. utriculariae* and *M. tetrahymenae* cells to co-evolve an adjusted physiology that reduces costs and increases benefits, driving the evolution of a stable endosymbiotic relationship.

## Materials and Methods

### *Tetrahymena* strains and culture conditions

*Tetrahymena utriculariae* was obtained from the laboratory of Dr. Thomas Posch at the University of Zurich (Switzerland). In 2005, *T. utriculariae* was discovered inside a trap of the aquatic carnivorous plant *U. reflexa* Oliv. collected by D. Sirová in the Okavango Delta, Botswana [13, 63]. *T. utriculariae* cells were grown in 2.5% (v/v) lettuce medium in modified Dryl’s solution [in which KH_2_PO_4_ was used instead of NaH_2_PO4_2_H_2_O] [64, 65] and fed with sufficient *Klebsiella pneumoniae* (NBRC 100048 strain) [66]. The *K. pneumoniae* cells were cultured overnight in Luria-Bertani (LB), added into the Dryl’s medium at 1000-fold dilution, incubated overnight, and then provided to the *T. utriculariae* cells. Cell cultures were maintained at 23 °C with a light-to-dark cycle of 12 h/12 h. The light was turned on at 8 AM and switched off at 8 PM. After sub-culturing, endosymbiotic *T. utricularia* cells reached early stationary phase on Day 5, whereas aposymbiotic cells took approximately 3 days to do so. All experiments were conducted using cells in early stationary phase.

To assess growth of *T. utriculariae* under both aerobic and hypoxic conditions, ∼1,000 cells were seeded in 2 ml of lettuce medium supplemented with *K. pneumoniae* (NBRC 100048 strain) for aerobic cultures. These cultures were maintained at room temperature (RT) with a 12 h/12 h light/dark cycle. For hypoxic cultures, the starting cell density was 1,000 cells/ml, and the cultures were maintained at 23 °C under hypoxic conditions with a 12 h/12 h light/dark cycle. Hypoxia was achieved by culturing the cells inside airtight glass vials, filling the vials completely with culture media containing *T. utriculariae* and without leaving any space for air. A parallel group of aerobic and hypoxic cultures was maintained under constant darkness to study the effect of light and photosynthesis by the endosymbiotic algae *M. tetrahymenae*. These dark-treated cultures allowed us to evaluate how the algae contribute to the growth of *T. utriculariae* under varying light conditions. Multiple tubes containing identical starter cultures (2 ml of the same cell density and hosting bacteria as food) were prepared to facilitate long-term growth analysis of hypoxic cultures. The hypoxic cultures were maintained for 70 days, with weekly cell counts. Three tubes were opened per condition for cell counting and they were discarded after use to avoid reintroduction into the hypoxic environment.

For aerobic cultures, 2 ml of fresh lettuce medium containing *K. pneumoniae* was added to each well on Day 3 to support growth. On Day 4, 1 ml of the same medium was added. On Day 5, 5 µL of *K. pneumoniae* grown in LB medium was added. To serve as controls for sufficient food supply, parallel cultures of *T. thermophila* were grown under identical aerobic conditions since they grew much faster and consumed more bacteria, including for the light/dark and constant-dark groups. This protocol facilitated a comparative evaluation of symbiotic and aposymbiotic *Tetrahymena* spp. under varying environmental and light conditions over an extended culture period.

### Obtaining a culture of free-living *Micractinium tetrahymenae*

*M. tetrahymenae* was isolated by plating endosymbiotic *T. utriculariae* on CA [67–69] agar plates. The culture plates were incubated at 23 °C with a 12/12 h light/dark cycle. Approximately two weeks later, a single colony was streaked on CA + ampicillin (100 mg/ml) agar plates, and another two weeks later, an axenic single colony was inoculated in CA + ampicillin (100 mg/ml) liquid culture.

### Genome sequencing, assembly, and annotation of *T. utriculariae*

After entering the early stationary phase, *T. utriculariae* cells were starved for an additional two days. Subsequently, they were collected by centrifugation at 1000 x *g* for 3 min. To extract high-quality total genomic DNA from the collected cells, ∼1×10^6^ cells were washed twice with modified Dryl’s solution [64]. The cells were then lysed with 0.25 M TCMS buffer and 0.3% (w/v) NP-40 in an equal volume. QIAGEN Genomic-tip 20/G (Qiagen, Germany) was used to extract the genomic DNA. The concentration of the extracted DNA was measured using a Qubit Fluorometer (Invitrogen, USA). Quality was checked using a Bioanalyzer 2100 system (Agilent Technologies, USA).

Long-read data was acquired from the PacBio Sequel platform (Pacific Biosciences, USA) and the Nanopore MinION platform (Oxford Nanopore Technologies, UK). A 20-kilobase (kb) insertion genomic DNA library was prepared according to a standard protocol and sequenced using a PacBio Sequel platform. Another genomic DNA library was prepared using a Ligation Sequencing kit (SQK-LSK 109, Oxford Nanopore Technologies) and sequenced on an Oxford Nanopore platform. For the short-read data, paired-end libraries were prepared using a standard protocol and sequenced using the Illumina Miseq and Nextseq platforms. All sequencing platforms were operated at the Genomics core of the Institute of Molecular Biology, Academia Sinica (https://www.imb.sinica.edu.tw/mdarray/).

The sequencing reads underwent quality trimming using Trimmomatic v0.38. The following options were used: ILLUMINACLIP=2:30:10, LEADING=3, TRAILING=3, SLIDINGWINDOW=4:15, and MINLEN=36. The subreads from the Pacbio platform and long reads over 1 kb with an average quality >q9 from the Nanopore sequencer were assembled using Canu v2.0. Assembly was performed with the following options: corMaxEvidenceErate=0.15 and genomeSize=100m. We successfully identified and assembled a total of 2089 contigs. To refine our data, we removed contigs that were either predicted to be a repeat or bubble of other contigs, circular in form, or shorter than 10 kb in length. After applying these filters, we were left with 518 contigs. From among these, we selected 216 contigs that had telomere repeats on at least one end for further assembly. We used Redundans v0.14a [70] to generate a haploid genome and to exclude contigs from two haplotypes of the same chromosomes during assembly. To do so, we applied the following criteria: alignment length should be >90% of the shorter contig, and sequence identity should be >90%. This process resulted in 174 chromosome-level contigs and 9 contigs with telomere repeats at one end. Next, we aligned the 174 chromosome-level contigs to the complete *T. thermophila* assembly and detected that contig 161 could align to two chromosomes in the assembly. We aligned the reads to contig 161 and observed that the reads connecting two homologous regions corresponding to the two *T. thermophila* chromosomes might be derived from the micronuclear genome. Therefore, we used Canu v2.0 to assemble the reads, which aligned to contig 161 and the nine contigs with telomere repeats on one end, using the options correctedErrorRate=0.15, corMaxEvidenceErate=0.15, and genomeSize=2m. This process resulted in seven new chromosome-level contigs. Ultimately, we were able to assemble 181 chromosome-level contigs. The sequence was then polished in NextPolish v1.3.1 using the Nanopore reads and in Pilon v1.23 using the Illumina reads, using three iterations. A training set was generated in Augustus v3.2.3 [71] using the gene annotation of *T. thermophila* (2014 version) acquired from the *Tetrahymena* genome database wiki (http://ciliate.org/index.php/) [26, 72]. The training set was further optimized by running optimize_augustus.pl. Then, we used BRAKER2 v2.1.4 [73] to annotate the genome, combining hints from the RNA-seq data and protein sequences of closely-related species (*T. thermophila*, *T. malaccensis*, *T. elliotti*, *T. borealis*) and the aforementioned training set with options AUGUSTUS_ab_initio, -- translation_table=6.

### Genome sequencing, assembly, and annotation of *M. tetrahymenae*

Genomic DNA extraction was performed using a modified CTAB-based method, adapted from Jagielski et al. [74]. The CTAB buffer was prepared using 1% CTAB, 1.4 M NaCl, 100 mM Tris-HCl (pH 8), 1% Triton X-100, and 20 mM EDTA (pH 8). TE buffer consisted of 10 mM Tris-HCl (pH 8) and 1 mM EDTA (pH 8). A phenol/chloroform/isoamyl alcohol mixture (25:24:1) with 0.1% 8-hydroxylquinoline (PCIA) was used, along with RNase A (10 mg/ml), 100% ethanol, 70% ethanol, and 3 M sodium acetate (NaOAc, pH 5.2). Approximately 109 *M. tetrahymenae* cells were harvested and washed twice with TE buffer by centrifugation at 10000 rcf for 30 seconds each time. The pellet was resuspended in 750 μl of CTAB extraction buffer and homogenized with an equal volume of 0.5 mm beads at 6 m/s for 80 seconds. The homogenate was then centrifuged at maximum speed for 5 minutes, and the aqueous phase was transferred to a new tube. Proteinase K (160 μg/ml) was added, and the mixture was incubated at 56 °C for 1 hour. Next, 500 μl of 5 M NaCl was added and thoroughly mixed, followed by the addition of 400 μl of prewarmed (65 °C) CTAB buffer, and the solution was incubated at 65 °C for 10 minutes. The lysate was processed three times with an equal volume of PCIA by thorough mixing and centrifugation at maximum speed for 5 minutes, transferring the aqueous phase to a fresh tube each time. DNA was precipitated by adding 1 ml of 100% ethanol and 20 μl of NaOAc, followed by incubation at RT for 30 minutes. The sample was centrifuged at 16000 rcf for 5 minutes, and the supernatant was discarded. The pellet was washed with 100 μl of 70% ethanol and centrifuged for 1 minute at 16000 rcf. The pellet was then air-dried for 10 minutes and rehydrated in 400 μl of TE buffer with 3 μl of RNase A, followed by incubation at 37 °C for 1 hour to remove RNA contamination. Subsequently, 1 ml of 100% ethanol and 20 μl of NaOAc were added to the suspension to precipitate the nucleic acids again at RT for 30 minutes. After centrifugation at 16000 rcf for 5 minutes, the supernatant was discarded, and the pellet was washed with 100 μl of 70% ethanol and centrifuged for 1 minute at 16000 rcf. Finally, the pellet was air-dried for 10 minutes and rehydrated in 50 μl of TE buffer. Genomic DNA integrity and quality were validated using a Bioanalyzer 2100 system (Agilent Technologies, USA).

The whole genome of *M. tetrahymenae* was sequenced using both Illumina paired-end (PE) sequencing and Oxford Nanopore Technologies (ONT). Illumina PE sequencing was utilized to generate high-accuracy short reads, whereas Nanopore sequencing provided long reads for comprehensive genome assembly. Quality control of the raw sequencing reads was conducted using NanoPlot [75] to evaluate read length and quality metrics. The data were assembled *de novo* using Canu v2.0 (minReadLength=5000 and genomeSize=65M) [76], which performed read correction, trimming, and assembly utilizing the best overlap graph (BOG) algorithm. The initial assembly was evaluated in QUAST [77] to assess metrics such as N50 and overall assembly completeness. The assembly was then refined through iterative polishing, first using Pilon [78] over three rounds with Illumina short reads, followed by three additional rounds of polishing with NextPolish [79], integrating both long and short reads for enhanced accuracy. Telomeric regions were identified using Tandem Repeat Finder (TRF) [80] alongside custom scripts for precise detection. Genome annotation was performed using BRAKER2 [73], incorporating RNA-seq data to improve gene prediction accuracy. GeneMark-ES [81], an *ab initio* gene predictor embedded within BRAKER2, was used to generate initial gene models. BUSCO [82] analysis was performed to evaluate genome completeness, utilizing single-copy orthologous genes as markers across the genome, transcriptome, and proteome datasets.

### Protein function prediction

We aimed to predict the functions of proteins encoded by the *T. utriculariae* and *M. tetrahymenae* genomes by means of multiple annotation approaches. First, we used InterProScan-5.40-77 [83] with the *goterms* option to assign Gene Ontology (GO) terms to proteins.

For *T. utriculariae*, we improved functional annotation by employing DScript 0.2.2 [84] with the trained model topsy_turvy_v1.sav, which estimates the probability of physical interactions between protein pairs. We set a stringent interaction probability threshold of 0.92, ensuring the density of predicted positive interactions (0.79%) aligned with that observed in *Saccharomyces cerevisiae* (0.78%) based on the BioGRID database [85]. For genes without functional annotations, we performed enrichment analysis on a gene-by-gene basis to determine if their interacting partners shared enriched GO terms. This analysis used Fisher’s exact test with Bonferroni correction, considering adjusted p-values below 1e-10 as statistically significant. Using this approach, we assigned functional predictions to 4,769 previously unannotated genes.

For *M. tetrahymenae*, we supplemented annotations using the MapMan Mercator4 v7.0 system [86]. Additionally, for genes lacking functional annotations in both *T. utriculariae* and *M. tetrahymenae*, we applied PANNZER2 [87], the OMA browser functional annotation tool [88], and STRING (version 12.0) [89] to further improve coverage.

As a result, we successfully annotated 20,777 out of 23,219 *T. utriculariae* genes and 13,630 out of 14,229 *M. tetrahymenae* genes with GO terms, thereby significantly enhancing functional annotation of both genomes.

### Synteny analysis

To analyze synteny between *T. thermophila* and *T. utriculariae*, first we performed an all-versus-all homology search using protein sequences from both species with DIAMOND v0.9.14 [90]. The DIAMOND search was conducted using an E-value threshold of 1e−10, restricting results to the top five hits per query, and outputs were formatted in m8 format. Subsequently, we utilized DupGen_finder (https://github.com/qiao-xin/DupGen_finder), an automated pipeline for detecting and classifying gene duplication events, integrating MCScanX [91] to identify collinear (syntenic) gene pairs between the two genomes. Whole-genome duplication (WGD)-derived gene pairs identified by MCScanX were excluded to specifically focus on non-WGD duplication modes. Remaining gene pairs were then classified based on genomic location, facilitating the identification of conserved genomic regions and providing insights into genome evolution and structural conservation between *T. thermophila* and *T. utriculariae*.

### Orthogroup identification

High-quality genome data for various *Tetrahymena* species (Supplementary Figure S1A) were sourced from the *Tetrahymena* genome database (https://tet.ciliate.org/) [72] and the P10K database (https://ngdc.cncb.ac.cn/p10k/) [92]. The freshwater obligate fish parasite *I. multifiliis* (https://ich.ciliate.org/), a close relative of *Tetrahymena* spp., [23] was selected as an outgroup. OrthoFinder (v2.5.5) [93] with a default inflation parameter of 1.5 was used to identify the orthologous genes among different ciliate species. Some of the orthogroups contained genes from only one species, and some genes that were not assigned to any orthogroups are defined as species-specific genes (Fig. 1C). Similar methods were used for *M. tetrahymenae* orthogroup prediction using the species mentioned in Figure 4A.

### Molecular phylogeny and divergence time analysis

Orthogroups that included proteins from all considered species, for both the *T. utriculariae* and *M. tetrahymenae* analyses, were concatenated and aligned using MAFFT (v7.525) [94], with trimming performed according to the default parameters of OrthoFinder. A maximum likelihood tree was constructed in RAxML-NG [95] using the concatenated orthogroup sequences and the LG+G4 sequence evolution model. A previous phylogenomic study based on the ciliate fossil record [96] indicated that *Ichthyophthirius* and *Tetrahymena* diverged ∼450 MYA [97, 98]. Accordingly, the divergence time of *T. utriculariae* from its close relatives was determined in the r8s package [99], with the maximum likelihood tree calibrated based on the divergence time of *Ichthyophthirius* and *Tetrahymena*.

We utilized CAFE 5 [100] to analyze gene family expansion and contraction, applying a random birth-and-death model on the divergence-time-aware tree constructed in r8s [99]. Additionally, we performed another round of OrthoFinder analysis using the three most closely related *Tetrahymena* species, i.e., *T. utriculariae*, *T. malaccensis*, and *T. thermophila*, with *T. elliotti* designated as an outgroup. Orthologous genes from these closely related species were employed for the gene family expansion and contraction analysis.

### Amino acid identity analysis

To determine pairwise amino acid identity (AAI) values between protein sequences in our genome dataset, we utilized the CompareM v0.1.2 program [101] (https://github.com/dparks1134/CompareM). We applied default settings and included the ‘--proteins --cpus 32 --keep_rbhs’ flags (with defaults of --evalue 1e-5, --per_identity 0.3, --per_aln_len 0.7). Additionally, we employed Reciprocal BLAST Hits (RBH) cross-matching to identify RBHs across all species pairs. We calculated AAI values by utilizing a consistent set of 9430 proteins for each species.

### Gene ontology enrichment

We built a customized org.db package for *T. utriculariae* and *M. tetrahymenae* gene ontology (GO) information, utilizing the makeOrgPackage function from the AnnotationForge [102] package provided by Bioconductor [103]. All GO term enrichment analyses were conducted using Clusterprofiler (v 4.10.1), and GO term redundancy was reduced using REVIGO [104] with default parameters. The reduced set of GO terms was further selected based on enrichment scores.

### Organellar genome assembly and annotation

For the mitochondrial genome, initially we employed the mitochondrial sequence of *T. thermophila* as a query in a BLAST search against the *T. utriculariae* contigs obtained from the Canu assembly, which did not include contigs with telomeres. Subsequently, we extracted the contigs that exhibited a match with the *T. thermophila* mitochondrial sequence. Using reads that could be mapped to these contigs, we performed a secondary assembly in Canu. Then, we applied a polishing step following the same process as employed for the nuclear genome. We employed Augustus to annotate the mitochondrial genome, and the hint file was generated through Exonerate with the options -m protein2genome --maxintron 500 --geneticcode 4, using the protein sequence encoded by the *T. thermophila* mitochondria. Following a methodology analogous to that applied to the nuclear genome, we compiled a training set based on the gene annotations of the *T. thermophila* mitochondria for subsequent use in the Augustus annotation process. Once we had obtained the mitochondrial genome fasta and CDS sequence files from Augustus, we further annotated and refined the final mitochondrial genome using GeSeq [105], MFannot [106], and Infernal (v1.1.5) [107]. GeSeq, Mfannot, and Infernal were also used to compare the other *Tetrahymena* mitogenomes, as presented in Supporting Table 2.

For nuclear-encoded mitogen annotation, we utilized the *T. thermophila* mitochondrial proteins isolated and reported by Smith et al. [108] to perform a blastp search against *T. utriculariae* proteins (with an e-value < 0.00001) to identify *T. utriculariae* mitochondrial proteins. Subsequently, we employed the orthologous gene (OG) tree of these identified proteins (513 OGs) as input to construct the species tree by means of STAG (Species Tree from All Genes) analysis.

For *M. tetrahymenae*, the mitochondrial and chloroplast genomes were assembled in a similar way to that mentioned above for *T. utriculariae*, using the reference mitochondrial and chloroplast genomes listed in Supplementary Tables S3 and S4. However, only GeSeq was used to annotate both the *M. tetrahymenae* mitochondrial and chloroplast genomes [105].

### Transmission Electron Microscopy (TEM) imaging

TEM sample preparation was conducted according to a microwave-assisted fixation protocol to examine mitochondrial ultrastructure and perialgal vacuole interactions in symbiotic *T. utriculariae*. Cells were fixed in 2.5% glutaraldehyde and 4% paraformaldehyde in 100 mM sodium cacodylate buffer (pH 7.2) using eight cycles of microwave irradiation (500W, 1 min) under vacuum, followed by room temperature incubation and overnight storage at 4 °C. After buffer rinses, samples underwent post-fixation with 1% osmium tetroxide using similar microwave cycling (450W). Samples were then water-rinsed and stained *en bloc* with 2% uranyl acetate under microwave irradiation. Dehydration proceeded through an ethanol gradient (10-100%) with brief microwave treatments at each concentration. Infiltration used Spurr’s resin beginning at 5% concentration, gradually increasing to 100% with vacuuming and regular agitation, followed by polymerization at 70 °C for 24 hours. Ultrathin sections (60-90 nm) were collected on copper grids, stained with lead citrate, and imaged using a transmission electron microscope at 100 kV. Images were analyzed for mitochondrial morphology, perialgal vacuole associations, and ultrastructural differences between symbiotic and aposymbiotic cells.

### Etomoxir treatment assay

To investigate the role of mitochondrial fatty acid oxidation in *T. utriculariae*, we performed pharmacological inhibition using etomoxir (ETO), an irreversible inhibitor of carnitine palmitoyltransferase 1 (CAS: 124083-20-1, MedChemExpress). Cultures were established in 1 mL microtiter plates with an initial concentration of 10,000 cells/mL in 2.5% (v/v) lettuce medium in modified Dryl’s solution, supplemented with *K. pneumoniae* (NBRC 100048) as a food source. We maintained cultures at 23 °C under a 12/12 h light/dark cycle without agitation. Experimental groups received final concentrations of 0 µM (control with equivalent ddH_2_O volume), 25 µM, or 50 µM ETO. Cell growth was monitored every 3 days by counting live cells under a microscope, with at least three biological replicates per condition to ensure reproducibility. For endosymbiont quantification, we employed a slide-pressing method, whereby appropriately diluted cells were placed on glass slides and gently pressed under a coverslip to lyse the host cells, enabling clear visualization and accurate counting of released *M. tetrahymenae* endosymbionts. We utilized fluorescence microscopy (Axio Observer, Carl Zeiss) to determine chlorophyll autofluorescence, analyzing at least 50 host cells per condition for statistically robust quantification.

### Transcriptome sequencing and analysis

#### Tetrahymena utriculariae RNA isolation

RNA was extracted from both symbiotic and aposymbiotic cells in stationary phase using a combination of TRI reagent (Sigma, Merck, USA) and a Qiagen RNeasy mini kit (Qiagen, Germany). The cells were pelleted by centrifuging at 1000 x *g* for 3 minutes, after which the pellet was washed twice with modified Dryl’s solution. The cells were then resuspended in 200 µl nuclease-free water. To lyse the cells, 1 ml of TRI reagent was added to the suspended cells, and the mixture was gently pipetted to ensure proper homogenization. After incubating for a further 5 minutes at RT, 50 mM of potassium acetate was added to the mixture and incubated again for 5 minutes at room temperature. The algal endosymbionts and debris were separated from the rest of the mixture by centrifuging at 12000 x *g* for 10 minutes at 4 °C. The resulting supernatant was moved to another clean 2 ml tube, before adding 300 µl of chloroform. The mixture was mixed vigorously for 3 minutes and then centrifuged at 21000 x *g* for at least 20 minutes at 4 °C to separate the organic and aqueous phases. The upper aqueous phase was mixed with an equal volume of 100% ice-cold ethanol. The mixture was remixed by pipetting and transferred to Qiagen RNeasy columns. RNA extraction was performed according to the column manufacturer’s instructions. RNA yield and purity were measured using a Qubit Fluorometer (Invitrogen, USA), and RNA integrity and quality were checked using a Bioanalyzer 2100 system (Agilent Technologies, USA).

#### Micractinium tetrahymenae RNA isolation

Endosymbiotic *M. tetrahymena* were isolated from symbiotic early stationary phase *T. utriculariae* cells using a sonicator. The cells were washed five times with nuclease-free water. Free-living *M. tetrahymenae* at log phase were used as a control. Total mRNA was extracted according to the same method as described above for *T. utriculariae*, with an additional step to break algal cells using 0.5 ml of 0.5 mm glass beads. Cells were beaten after adding 1 ml TRI reagent using a MP-Bio Fastprep-24 G bead beater at 6 ms^-1^ for 80 seconds, followed by 12000 x *g* centrifugation for 10 minutes at 4 °C. The supernatant was transferred to a new tube and mixed vigorously for 3 minutes with 300 µl of chloroform. The subsequent steps were the same as those described above to extract *T. utriculariae* mRNA.

#### Library preparation and mRNA sequencing

To prepare the RNA libraries, we used the SureSelect Strand-Specific RNA Library Preparation kit for Illumina (Agilent Technologies, USA) in accordance with the manufacturer’s instructions. The libraries were sequenced using a NovaSeq 6000 (Illumina, USA) sequencing system at Welgene Biotech Co. Ltd. (Taiwan). We sequenced three biological replicates each from symbiotic and aposymbiotic cells, with 150-bp paired-end reads per replicate. Fastqc v0.11.8 [109] and fastp v0.24.0 [110] were used for quality control and to trim the raw data, respectively. We used kallisto v0.51.0 using default parameters for transcript quantification [111].

#### Differential gene expression analysis

Differential expression analyses were carried out separately for the host *T. utriculariae* and its endosymbiont *M. tetrahymenae* using the limma framework in R [112]. For each organism, raw values of transcripts per million reads (TPM) were first filtered to retain only genes with TPM ≥ 0.005 in at least 70% of samples. The filtered counts were then log_2_-transformed (with a small pseudocount added to avoid zeros). The sample metadata were assembled to define, for *T. utriculariae*, the interaction of cell type (symbiotic vs. aposymbiotic) and time-point, and for *M. tetrahymenae*, the endosymbiotic status (Endo vs. Exo).

Linear models were fitted to the log_2_-TPM matrix via lmFit, and empirical Bayes moderation (eBayes) was applied to stabilize per-gene variance estimates and calculate moderated t-statistics [113]. For each gene and contrast, raw p-values were obtained from the moderated t-distribution (with gene-specific, augmented degrees of freedom) and then adjusted using the Benjamini–Hochberg procedure to control the false discovery rate [114]. Genes exhibiting |log_2_ fold-change| ≥ 1 and an adjusted p-value ≤ 0.01 were classified as significantly up- or downregulated under the specified conditions.

#### K-means clustering and heatmap visualization of *T. utriculariae* DEGs

Differentially expressed genes (|log_2_ fold-change| ≥ 1, adjusted p ≤ 0.01) were clustered on their z-score–normalized expression profiles across the 12 samples (six symbiotic, six aposymbiotic). Clustering was performed using the KMeans_rcpp function in ClusterR [115], specifying k = 4 clusters with k-means**++** initialization, 30 random restarts (num_init = 30), and up to 300 iterations per run (max_iters = 300). The set.seed(123) function ensured reproducibility. The resulting cluster assignments were visualized as a heatmap in the ComplexHeatmap package [116, 117], ordering rows by each gene’s peak absolute expression and splitting columns by symbiotic status (symbiotic vs. aposymbiotic), with an overlaid annotation of light regime (dark vs. light).

#### Measuring chlorophyll content

Equal numbers of endosymbiotic and free-living *M. tetrahymenae* cells were harvested and pelleted (3,000 × *g*, 5 min, 4 °C), followed by extraction in 1 mL of 100% methanol, with cells either ground in liquid nitrogen or bead-beaten in the dark for 20 minutes at 4 °C. After clarification (14,000 × *g*, 5 min, 4 °C), absorbance was measured using a 1 cm path-length cuvette at 652 nm (A_652_) and 665 nm (A_665_). Chlorophyll concentrations were calculated using the equations: Chlorophyll a (µg/mL) = 16.29 × A_665_ − 8.54 × A_652_; Chlorophyll b (µg/mL) = 30.66 × A_652_ − 13.58 × A_665_, and total chlorophyll was determined as the sum of Chlorophyll a and Chlorophyll b [118, 119]. Measurements were conducted in triplicate, and statistical comparisons were performed by two-tailed t-tests.

## Data availability

The polished genome, gene annotation, CDS sequences, and protein sequences have all been uploaded to NCBI (*T. utriculariae*: PRJNA1236653, *M. tetrahymenae*: PRJNA1243065). Gene annotations and genomic data are available from the *Tetrahymena* Genome Database (tet.ciliate.org). Genome and RNA sequencing data from the current work have been deposited to the NCBI database (*T. utriculariae*: PRJNA1236653; *M. tetrahymenae*: PRJNA1243065).

## Acknowledgments

We thank members of the Leu lab for helpful discussion and comments on the manuscript. We also thank John O’Brien for manuscript editing, the IMB Genomics and Imaging Cores for experimental assistance. This work was supported by Academia Sinica of Taiwan (grant no. AS-IA-110-L01 and AS-GCS-113-L03) and the National Science and Technology Council of Taiwan (NSTC 113-2326-B-001-002). KMM was supported by an NSTC postdoctoral fellowship (NSTC 113-2811-B-001-065).

## Author contributions

JYL conceived the study. KMM, YHC, and JYL designed analyses and interpreted results. KMM, YHC, PTN and LWC performed the experiments. KMM, YHC, CFJL, and CWL performed data analysis. KMM, YHC, and JYL wrote the paper. TP provided the *Tetrahymena* strain. All authors read and approved the final manuscript.

## Supplementary figure

**Supplementary Figure S1:**
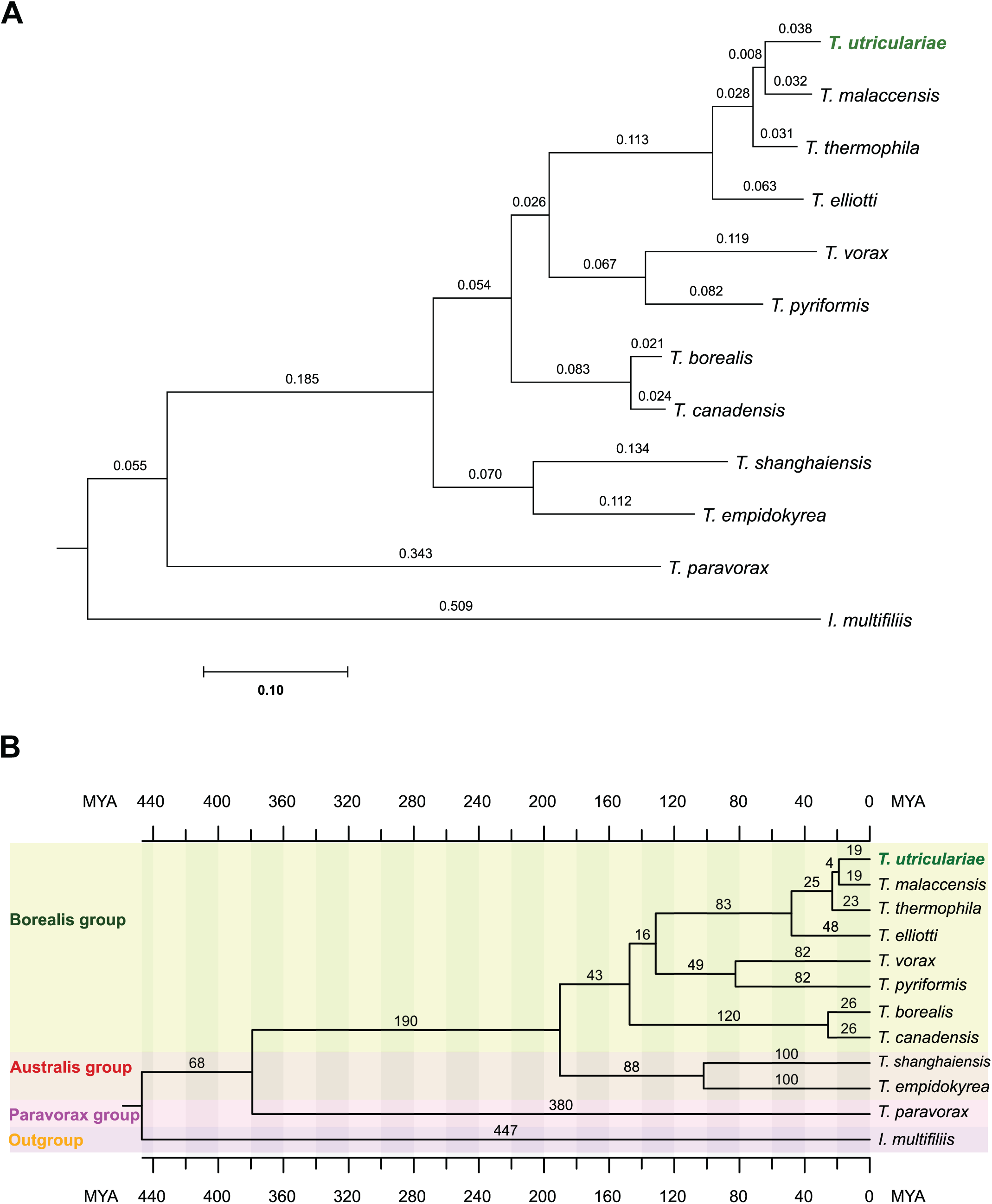
Phylogenetic relationships and evolutionary timeline of *Tetrahymena* species. (A) Maximum likelihood phylogenetic tree of 12 *Tetrahymena* species with *Ichthyophthirius multifiliis* as the outgroup. The tree was constructed from a concatenated alignment of single-copy orthologous genes identified by OrthoFinder. Branch lengths represent genetic distance (scale bar: 0.10 substitutions per site), and values at nodes indicate branch distances. *T. utriculariae* forms a well-supported clade with *T. malaccensis* and *T. thermophila*, with *T. utriculariae* being most closely related to *T. malaccensis*, followed by *T. thermophila*. (B) Time-calibrated phylogeny showing the estimated divergence times among *Tetrahymena* species in millions of years ago (MYA). The tree reveals four major phylogenetic groups: Borealis group (green, including *T. utriculariae*), Australis group (red), Paravorax group (pink), and the outgroup *I. multifiliis* (orange). The *Tetrahymena* genus originated approximately 447 MYA, with major diversification events occurring at 380 MYA (Paravorax divergence) and 190 MYA (Borealis-Australis split). Within the Borealis group, *T. utriculariae* diverged from *T. malaccensis* approximately 19 MYA, i.e., shortly after its divergence from *T. thermophila* (23 MYA). This recent divergence indicates that evolution of a symbiotic lifestyle in *T. utriculariae* is a relatively novel adaptation in the genus.

**Supplementary Figure S2:**
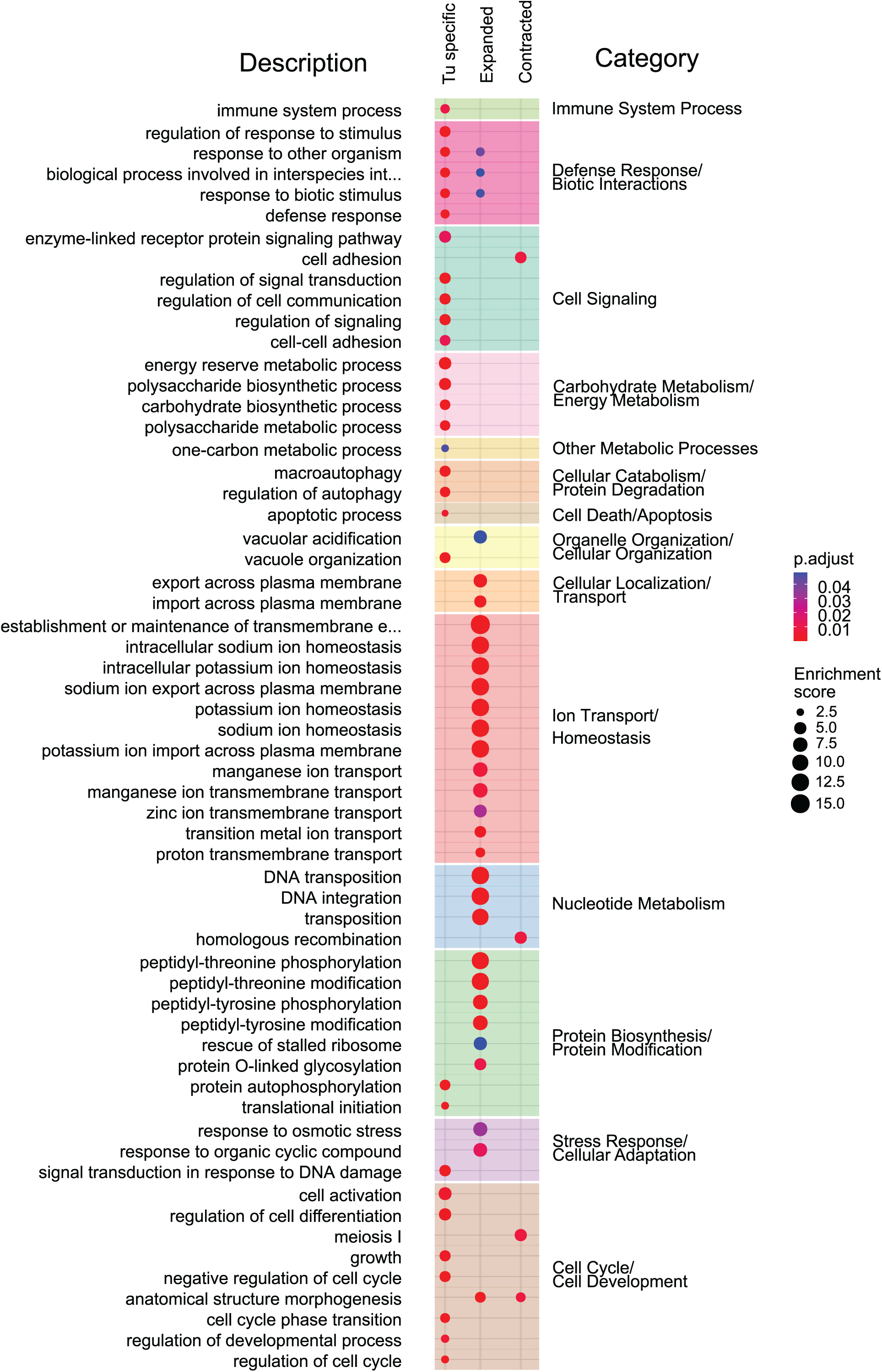
GO enrichment analysis of *T. utriculariae* gene categories: *T. utriculariae*-specific, expanded, and contracted genes. *T. utriculariae*-specific genes: Gene Ontology (GO) terms significantly enriched in genes unique to *T. utriculariae*, notably in ion transport, protein modification, and stress responses. Expanded genes: GO terms enriched in gene families that have expanded in *T. utriculariae*, especially those involved in ion homeostasis, signal transduction, and protein modification. Contracted genes: GO terms enriched in gene families that have contracted in *T. utriculariae*, particularly in developmental processes and cell adhesion.

**Supplementary Figure S3:**
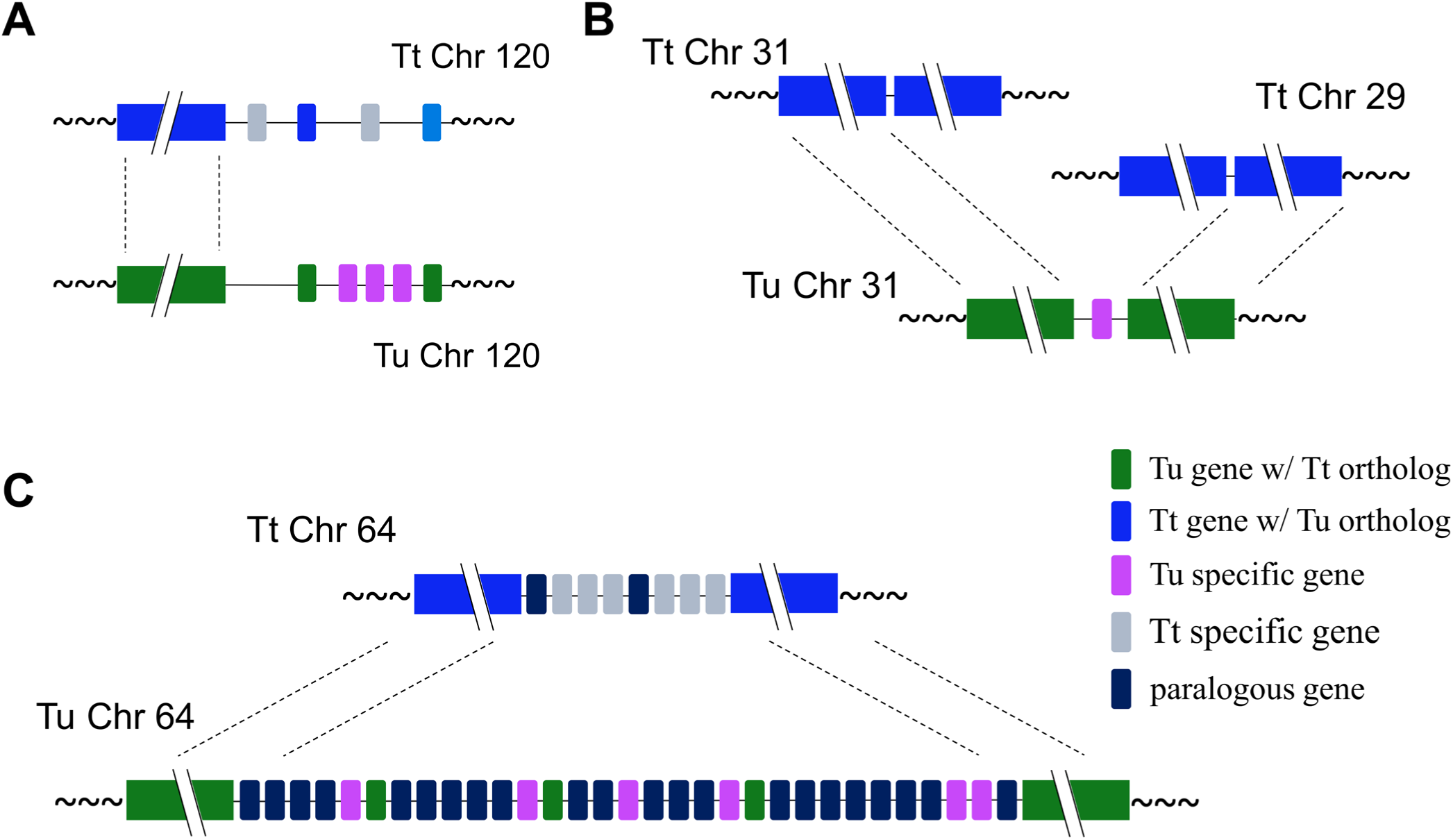
Comparative genomic architecture reveals chromosomal evolution between *T. utriculariae* and *T. thermophila*. (A) An example of telomeric species-specific genes. Comparative analysis of chromosome 120 between *T. utriculariae* (Tu) and *T. thermophila* (Tt) reveals a conserved central region with syntenic genes (green and blue blocks) and terminal regions with *T. utriculariae*-specific genes (purple blocks). (B) An example of species-specific genes in chromosomal fusion regions. A chromosomal rearrangement where genetic material from two separate *T. thermophila* chromosomes (Tt Chr 31 and Tt Chr 29) has been consolidated into a single chromosome in *T. utriculariae* (Tu Chr 31). Syntenic blocks are shown in green and blue, whereas non-syntenic species-specific genes are highlighted in purple. (C) An example of species-specific genes in expanded chromosomal regions. *T. utriculariae* chromosome 64 exhibits a substantial length increase compared to its *T. thermophila* counterpart. Paralogous genes resulting from duplication events are shown in dark blue blocks, representing a mechanism of genomic expansion in *T. utriculariae*.

**Supplementary Figure S4:**
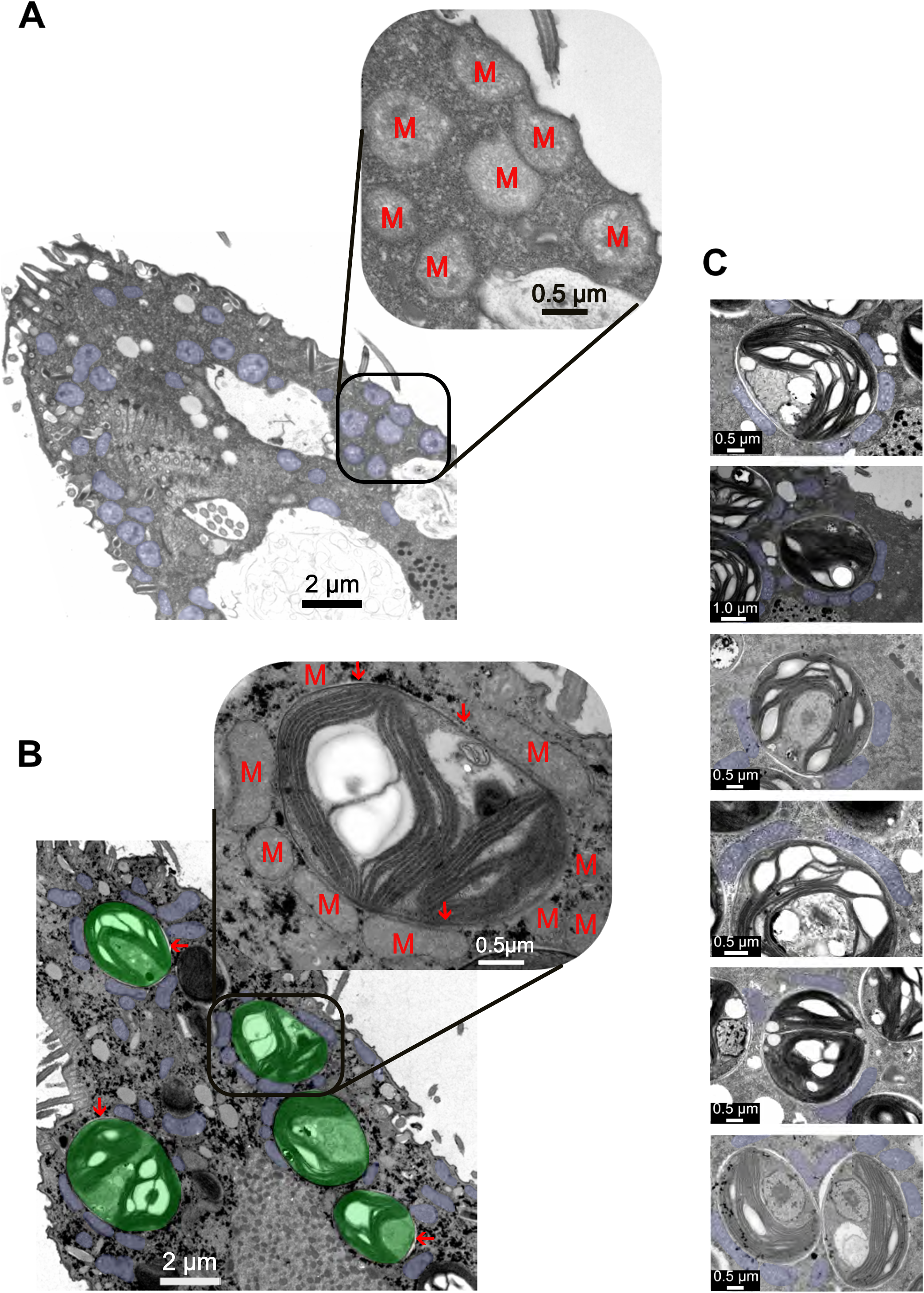
Distinct mitochondrial morphologies in aposymbiotic versus symbiotic *T. utriculariae* cells. Continuation from Figure 2D. (A) Transmission electron micrograph of an aposymbiotic *T. utriculariae* cell with mitochondria highlighted in blue pseudocolor. In the absence of endosymbionts, mitochondria display predominantly rounded to oval morphologies distributed throughout the cytoplasm. High magnification inset shows the typical spherical mitochondrial ultrastructure with clearly visible cristae (scale bar: 0.5 μm). The main image scale bar represents 2 μm. (B) Transmission electron micrograph of a symbiotic *T. utriculariae* cell containing multiple *M. tetrahymenae* endosymbionts (pseudocolored green). Mitochondria (blue pseudocolor) show dramatically altered morphology compared to aposymbiotic cells, being elongated and closely associated with the perialgal vacuole membrane surrounding the endosymbionts. High magnification inset (scale bar: 0.5 μm) reveals the intimate physical association between host mitochondria (M) and the perialgal vacuole membrane (red arrow), with mitochondria forming extensive membrane contacts and wrapping around the perialgal vacuole membrane (PVM) boundary. Note that mitochondria not associated with endosymbiont-containing vacuoles retain a more conventional morphology. Main image scale bar: 2 μm. (C) Gallery of additional symbiotic *T. utriculariae* cells maintained under low oxygen conditions, showing consistent mitochondria-per algal vacuole (PV) associations. Each panel demonstrates the characteristic elongation and wrapping of mitochondria (blue pseudocolor) around the endosymbiotic algae contained within the perialgal vacuole membrane (PVM). This structural adaptation likely facilitates metabolic exchange between host and endosymbiont, particularly under hypoxic conditions where both partners may engage in complementary energy metabolism. Scale bars: 0.5 μm.

**Supplementary Figure S5:**
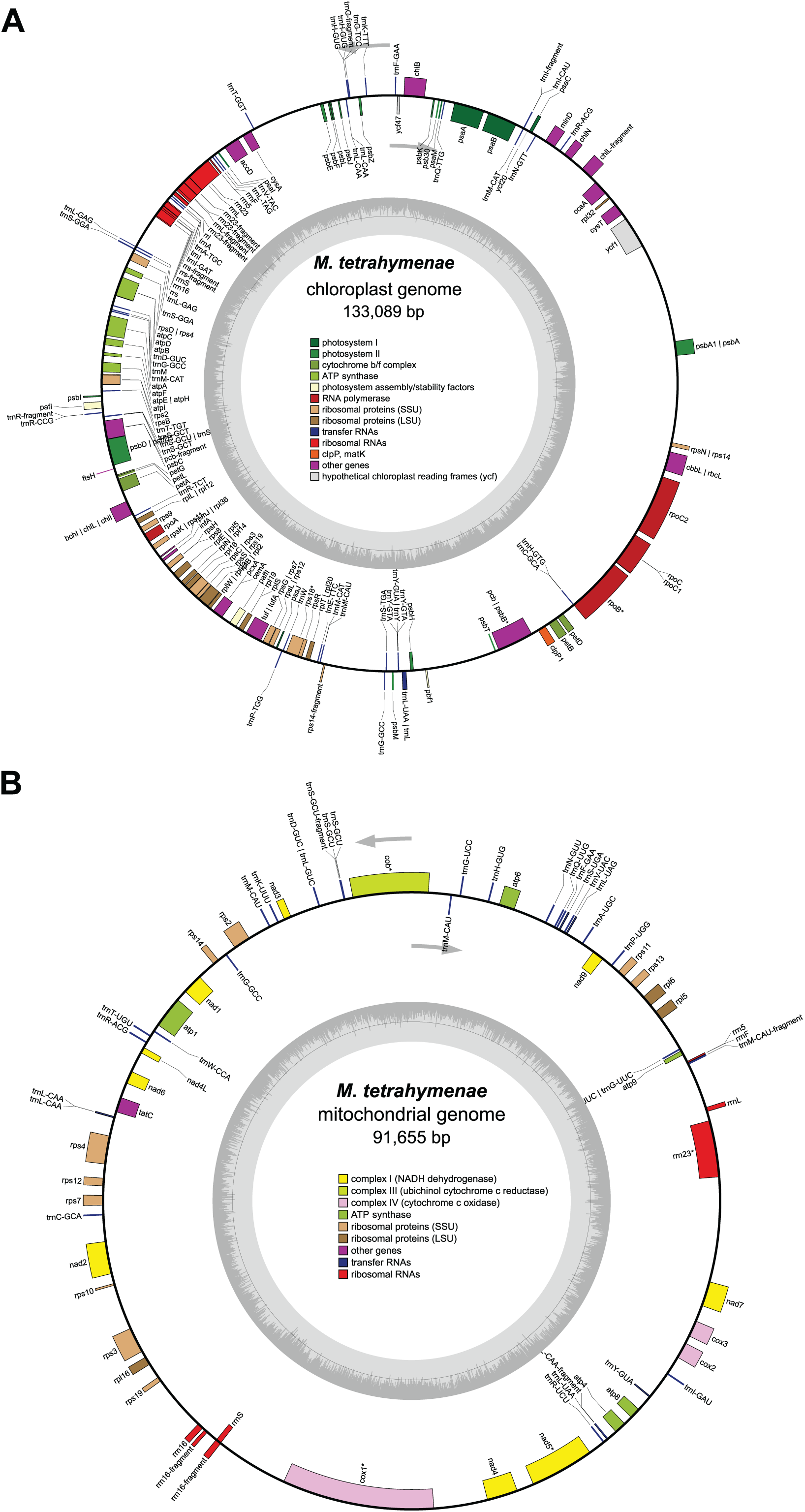
Organellar genome architecture of the endosymbiotic alga *M. tetrahymenae*. (A) Chloroplast genome map of *M. tetrahymenae*. The circular chloroplast genome (133,089 bp) contains a typical complement of photosynthetic and housekeeping genes. Genes are color-coded by function as indicated in the legend: photosystem I (dark green), photosystem II (light green), cytochrome b/f complex (olive), ATP synthase (yellow), photosystem assembly/stability factors (orange), RNA polymerase (red), ribosomal proteins (brown; SSU: small subunit, LSU: large subunit), transfer RNAs (blue), ribosomal RNAs (orange-red), open reading frames (ORFs, purple), other genes (white), and hypothetical chloroplast reading frames (gray). Gene names are labeled on the outside of the circle, with genes on the outer strand shown on the outside and genes on the inner strand shown on the inside. The inner gray circle represents GC content variation across the genome. (B) Mitochondrial genome map of *M. tetrahymenae*. The circular mitochondrial genome (91,655 bp) encodes proteins involved in cellular respiration and genetic information processing. Genes are color-coded by function: complex I/NADH dehydrogenase (yellow), complex III/ubiquinol-cytochrome c reductase (orange), complex IV/cytochrome c oxidase (pink), ATP synthase (green), ribosomal proteins (brown; SSU: small subunit, LSU: large subunit), other genes (purple), transfer RNAs (blue), and ribosomal RNAs (red). Gene names are labeled on the outer circle, with orientation indicating the strand location. The inner gray circle depicts GC content across the genome. Gray arrows indicate the location of large repeat regions within the mitochondrial genome.

## Supplementary Tables

**Table S1.** Distribution of syntenic and non-syntenic genes in *T. utriculariae*.

|  | Genes in syntenic blocks | Genes not in syntenic blocks | Total |
| --- | --- | --- | --- |
| *T. utriculariae*-specific genes | 1,792 | 716 | 2,508 |
| Non-specific genes | 19,651 | 1,060 | 20,711 |
| **Total** | **21,443** | **1,776** | **23,219** |

**Table S2.** Mitochondrial genome comparison among *Tetrahymena* species that were used in phylogenetic analysis (Figure 2B and C).

| Species | *T. utriculariae* | *T. thermophila* | *T. malaccensis* | *T. paravorax* | *T. pyriformis* | *T. rostrata* |
| --- | --- | --- | --- | --- | --- | --- |
| **Genome Size (bp)** | 51725 | 47577 | 47691 | 47496 | 47296 | 47310 |
| **AT Content** | 78.53 | 79.24 | 80.09 | 81.53 | 78.67 | 78.29 |
| **Total Genes** | 58 | 60 | 59 | 60 | 58 | 59 |
| **Protein Coding Genes** | 44 | 44 | 45 | 44 | 45 | 45 |
| **tRNA Genes** | 7 | 8 | 8 | 10 | 8 | 8 |
| **rRNA Genes** | 6 | 7 | 6 | 6 | 6 | 6 |
| **Gene Density** | 1121.31 | 1261.11 | 1237.13 | 1263.26 | 1226.32 | 1247.09 |
| **Gene Proportion** | 87.67 | 95.91 | 95.46 | 94.62 | 95.75 | 95.82 |
| **Intergenic Proportion** | 12.63 | 4.09 | 4.54 | 5.65 | 4.25 | 4.18 |
| **Total Intergenic Length** | 6534 | 1947 | 2163 | 2684 | 2011 | 1978 |

**Table S3:** List of mitochondrial reference genomes used to assemble the *M. tetrahymenae* mitochondrial genome.

| Species | Strain/isolate |
| --- | --- |
| *Micractinium sp* | LBA 32 |
| *Chlorella sorokiniana* | 1230 |
| *Chlorella variabilis* | NC64 |
| *Chlorella vulgaris* | 211-11P |
| *Chlorella vulgaris* | NJ-7 |

**Table S4:**
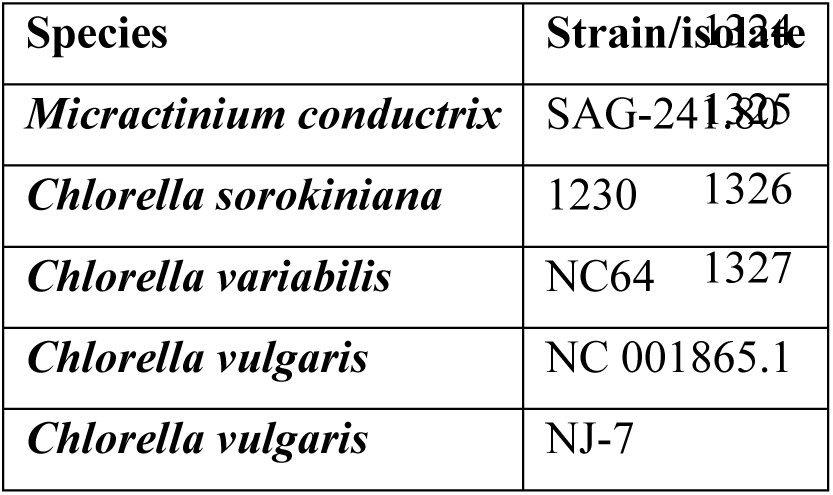
List of chloroplast reference genomes used to assemble the *M. tetrahymenae* chloroplast genome.

## Supplementary Data

**Supplementary Data S1: Orthogroup Analysis of *Tetrahymena* Species**

**S1A. T. utriculariae gene info**

Detailed annotation of *T. utriculariae* genes, including gene IDs, descriptions, symbols, UniProt IDs, and mini chromosome IDs.

**S1B. Shared by all three *Tetrahymena* species**

Orthologous gene clusters identified through OrthoFinder shared among *Tetrahymena utriculariae*, *T. malaccensis*, and *T. thermophila*.

**S1C. Specific to *T. utriculariae***

Genes exclusively identified in the *T. utriculariae* genome, annotated with gene IDs and descriptions.

**Supplementary Data S2: Genome-wide Gene Ontology (GO) and Gene Family Analysis of *Tetrahymena utriculariae***

**S2A. *T. utriculariae*-specific GO**

Gene Ontology enrichment analysis of *T. utriculariae*-specific genes.

**S2B. *T. utriculariae* expanded gene families**

Expanded gene families identified in *T. utriculariae* via CAFE5 analysis.

**S2C. *T. utriculariae* contracted gene families**

Contracted gene families identified in *T. utriculariae* via CAFE5 analysis.

**S2D. Expanded GO**

Gene ontology enrichment analysis for expanded gene families in *T. utriculariae*.

**S2E. Contracted GO**

Gene ontology enrichment analysis for contracted gene families in *T. utriculariae*.

**Supplementary Data S3: Transcriptomic Analysis of *Tetrahymena utriculariae***

**Metadata**

Metadata information describing RNA-seq samples, conditions, biological replicates, symbiotic states, and time-points.

**S3A. TPM**

Transcripts per million (TPM) data for *T. utriculariae* genes in symbiotic and aposymbiotic states across multiple diurnal time-points with biological replicates.

**S3B. Differentially expressed genes**

Differentially expressed genes (DEGs) between symbiotic and aposymbiotic conditions, including statistical parameters (fold-change, adjusted p-values).

**S3C. K-means clustering**

Expression profiles of DEGs grouped by k-means clustering into different temporal and condition-specific clusters.

**S3D. *T. utriculariae* gene ontology**

Gene ontology enrichment analyses for each DEG cluster, detailing functional implications in symbiotic versus aposymbiotic states.

**Supplementary Data S4: Transcriptomic Analysis of *Micractinium tetrahymenae***

***M. tetrahymenae* gene information**

Annotation details for *M. tetrahymenae* genes, including gene IDs, descriptions, standardized symbols, and UniProt IDs.

**S4A. M. tetrahymenae TPM**

Transcripts per million (TPM) data comparing endosymbiotic and free-living *M. tetrahymenae* conditions.

**S4B. *M. tetrahymenae* differentially expressed genes**

Differentially expressed genes (DEGs) between endosymbiotic and free-living conditions, with expression statistics and significance.

**S4C. *M. tetrahymenae* gene ontology**

Gene ontology enrichment analysis highlighting key metabolic and physiological shifts in endosymbiotic *M. tetrahymenae* compared to the free-living state.

